# Molecular recognition of two endogenous hormones by the human parathyroid hormone receptor-1

**DOI:** 10.1101/2022.10.09.511452

**Authors:** Li-Hua Zhao, Qingning Yuan, Antao Dai, Chuan-Wei Chen, Xinheng He, Chao Zhang, Youwei Xu, Yan Zhou, Ming-Wei Wang, Dehua Yang, H. Eric Xu

**Author notes:** Correspondence (Li-Hua Zhao); (Dehua Yang); (H. Eric Xu). These authors contributed equally: Li-Hua Zhao, Qingning Yuan and Antao Dai.

## Abstract

Parathyroid hormone (PTH) and PTH-related peptide (PTHrP) are two endogenous hormones recognized by PTH receptor −1 (PTH1R), a member of class B G protein couple receptors (GPCRs). Both PTH and PTHrP analogs including teriparatide and abaloparatide, are approved drugs for osteoporosis, but they exhibit distinct pharmacology. Here we report two cryo-EM structures of human PTH1R bound to PTH and PTHrP in the G protein-bound state at resolutions of 2.6 and 3.2 Å, respectively. Detailed analysis of these structures uncovers both common and unique features for the agonism of PTH and PTHrP. Molecular dynamics (MD) simulation and together with site-directed mutagenesis studies reveal the molecular basis of endogenous hormones recognition specificity and selectivity to PTH1R. These results provide a rational template for the clinical use of PTH and PTHrP analogs as an anabolic therapy for osteoporosis and other disorders.

## INTRODUCTION

Parathyroid hormone (PTH) and PTH-related protein (PTHrP) are two endogenous peptide hormones that play key and distinct biological roles in skeletal development, calcium- and phosphate-regulating actions, and bone turn over^1,2^. Lack of functional PTH results in hypoparathyroidism^3^, while overexpression of PTHrP derived from tumor cells is the most common reason for cancer-associated hypercalcemia^4^. Both PTH and PTHrP also have potent cardiovascular effects, including stimulating heart rate increase independent of autonomic reflexes^5^. Both PTH and PTHrP exert their effects on bone by activating the PTH type 1 receptor (PTH1R), primarily via G_s_-cAMP signaling^1,6^. High-resolution structures of PTH1R-G_s_ complexes bound to these endogenous hormones are important for both pharmacological research as well as clinical development aiming at PTH1R.

Osteoporosis is a disease of decreased bone mass, microarchitectural deterioration, and fragility fractures. It is widespread and can affect all ethnic groups. Osteoporosis becomes a major clinical problem in older women and men, especially in postmenopausal women^7^. Clinical trials were proved that both PTH and PTHrP can stimulate bone anabolism in patients with osteoporosis^6,8^. Teriparatide, recombinant human parathyroid hormone encompassing residues 1-34 (PTH1-34), and abaloparatide, recombinant human PTH-related protein encompassing residues 1-34 (PTHrP1-34), are two synthetic analogs of PTH and PTHrP. Although they exhibit distinct pharmacology, they were the only two approved drugs by FDA for the treatment of osteoporosis^6^. Recent studies suggest that PTH and PTHrP differ in their relative capacities to bind to two pharmacologically distinguishable high-affinity PTH1R conformations, R^0^ (the G protein-uncoupled PTH1R conformation) and RG (the G protein-coupled PTH1R conformation)^9,10^. PTH (1-34) can bind to the R^0^ conformation with the higher affinity than PTHrP (1-36) and prolonged cAMP responses^10^. Whereas PTHrP (1-36) binds weakly to R^0^, but it preferentially binds to the RG, promoted by the over expression of a high affinity of Gα_s_ and produce shorter cAMP responses^6,10^. Hattersley et al.^11^ found that abaloparatide has similar high affinity to the RG as teriparatide, and abaloparatide shows more selectivity for the RG and a shorter cAMP signaling response compared to teriparatide. Abaloparatide and teriparatide different selectivity for the RG conformation can explain the different increase in bone mineral density (BMD) and different incidence of hypercalcemia in the clinical trials treated by abaloparatide and teriparatide^6^. Such conformational selectivity of receptor binding can be a key regulator of that ligand’s biological activity, which provides an important clue to optimize conformational selective PTH and PTHrP analogs for a new treatment option for PTH-related disorders^12^.

Up to now, the most effective treatments of osteoporosis are based on daily injections teriparatide or abaloparatide. These therapies are both costly and inconvenient in clinical application due to the therapy by injection. However, there are no available small-molecule agonist drugs for the treatment of osteoporosis, in part due to lack of detailed structure information of an intact endogenous peptide-PTH1R-G-protein complex^13,14^. The only available structure of the full-length PTH1R is in complex with a synthetic long-acting PTH (LA-PTH), which reveals extensive interactions of LA-PTH with PTH1R^2^. The detailed structural and mechanistic understanding of the receptor activation by endogenous peptide hormones is critical for explaining how PTH and PTHrP mediate their distinct biologic actions via a common receptor and for aiding the rational development of new drug therapies targeting PTH1R. Therefore, we determined two cryo-electron microscopy (cryo-EM) structures of the active PTH1R– G_s_ complexes bound to PTH and PTHrP to provide insights into the pharmacological mechanisms for the clinical use of endogenous and clinical ligands.

## Materials and Methods

### Constructs

The human PTH1R (residues 27-502) with G188A and K484R mutations was cloned into pFastBac vector (Invitrogen) with the haemagglutinin signal peptide (HA) followed by a TEV protease cleavage site and a double MBP (2MBP) tag to facilitate expression and purification^2^. To facilitate a stable complex, the above PTH1R construct was added the LgBiT subunit (Promega) at the C terminus of PTH1R. A bovine dominant-negative human Gα_s_ (DNGα_s_) with mutations G226A and A366S, is a chimera that the AHD of Gα_s_ (69-205) was replaced with Gα_i_ (62-182) to bind to the Fab_G50^2^. Based on the published DNGα_s_, a modified bovine Gα_s_ (mDNGα_s_), its N terminus (M1–K25) and α-helical domain (AHD, F68–L203) of Gα_s_ were replaced by the N terminus (M1–M18) and AHD (Y61–K180) of the human Gα_i_, which can bind scFv16 and Fab_G50^18^ and the residues N254-T263 of Gα_s_ were deleted. In addition, eight mutations (G49D, E50N, L63Y, A249D, S252D, L272D, I372A, and V375I). To facilitate the folding of the G protein, DNGα_s_ and mDNGα_s_ was co-expressed with GST-Ric-8B^19^. Rat Gβ1 with an N-terminal MHHHHHHSSGLVPRGSHMASHHHHHHHHHH-tag (His16) was fused with the SmBiT subunit (peptide 86, Promega)^20^ via a 15 amino acid (GSSGGGGSGGGGSSG) linker at its C terminus. In addition, to clone the constructs into the pcDNA3.1 vector (Promega) for cAMP accumulation, all constructs were cloned using homologous recombination (Clone Express One Step Cloning Kit, Vazyme).

### Protein expression

The PTH1R and G proteins were co-expressed in *Sf9* insect cells (Invitrogen). When the cells grew to a density of 3.5 × 10^6^ cells/mL in ESF 921 cell culture medium (Expression Systems), we infected the cells with five separate virus preparations at a ratio of 1:2:2:2:2 for PTH1R(27-502)-LgBiT-2MBP, DNGα_s_ or mDNGα_s_, His16-Gβ1-peptide 86, Gγ2, and GST-Ric-8B. The infected cells were cultured at 27°C for 48 h before collection by centrifugation and the cell pellets were stored at −80°C.

### Expression and purification of Nb35

Nanobody-35 (Nb35) was expressed in *E. coli* BL21 cells, the cultured cells were grown in 2TB media with 100 μg/mL ampicillin, 2 mM MgCl_2_, 0.1% glucose at 37°C for 2.5 h until OD600 of 0.7-1.2 was reached. Then the culture was induced with 1 mM IPTG at 37°C for 4-5 h, and harvested and frozen at −80°C until use. Nb35 was purified by nickel affinity chromatography and followed by size-exclusion chromatography using HiLoad 16/600 Superdex 75 column following overnight dialysis against 20 mM HEPES, pH 7.4, 100 mM NaCl, 10% glycerol. The quality of purified protein was verified by SDS-PAGE and store at −80 °C.

### Complex purification

The complexes were purified according to previously described methods^21,22^. It was resuspended in 20 mM HEPES pH 7.4, 100 mM NaCl, 10 mM MgCl_2_, 10 mM CaCl_2_, 2 mM MnCl_2_, 10% glycerol, 0.1 mM TCEP, 15 μg/mL Nanobody (Nb35), 25 mU/mL apyrase (Sigma), 12 µM PTH1-34 (Synpeptide Co., Ltd) and 12 µM PTHrP1-36 (TGpeptide Biotechnology Co., Ltd), supplemented with Protease Inhibitor Cocktail (TargetMol, 1 mL/100 mL suspension). The lysate was incubated for 1 h at room temperature and complex from membranes solubilized by 0.5% (w/v) lauryl maltose neopentylglycol (LMNG, Anatrace) supplemented with 0.1% (w/v) cholesteryl hemisuccinate TRIS salt (CHS, Anatrace) for 2 h at 4°C. The supernatant was isolated by centrifugation at 65,000 × g for 40 min, and the solubilized complex was incubated with Amylose resin (NEB) for 2 h at 4°C. The resin was loaded onto a plastic gravity flow column and washed with 15 column volumes of 20 mM HEPES, pH 7.4, 100 mM NaCl,10% glycerol,10 mM MgCl_2_, 1 mM MnCl_2_, 0.01% (w/v) LMNG, 0.01% glyco-diosgenin (GDN, Anatrace) and 0.004% (w/v) CHS, 2 µM PTH1-34 or 2 µM PTHrP1-36, and 25 μM TCEP. After washing, the protein was treated overnight with TEV protease on column at 4°C. Next day the flow through was collected and concentrated, then UCN1-CRF2R-G_o_ and UCN1-CRF2R-G_11_ were loaded onto a Superdex200 10/300 GL column and Superose6 Increase 10/300GL (GE Healthcare), respectively, with the buffer containing 20 mM HEPES, pH 7.4, 100 mM NaCl, 2 mM MgCl_2_, 0.00075% (w/v) LMNG, 0.00025% GDN, 0.0002% (w/v) CHS, 2 µM PTH1-34 or 2 µM PTHrP1-36, and 100 μM TCEP. The complex fractions were collected and concentrated individually for electron microscopy experiments.

### Cryo-EM data acquisition

For the preparation of cryo-EM grids, 2.5 μL of the purified PTH-PTH1R-G_s_ and PTHrP-PTH1R-G_s_ complexes at a concentration of ∼8.4 mg/mL and ∼5.1 mg/mL were <u>respectively</u> applied to the glow-discharged Au 300 mesh holey carbon grids (Quantifoil R1.2/1.3). The grids were blotted and then plunge-frozen in liquid ethane using a Vitrobot Mark IV (ThermoFisher Scientific) at 4°C.

Cryo-EM images of PTH-PTH1R-G_s_ were collected on a Titan Krios G4 equipped with a Gatan K3 direct electron detector with super-resolution mode and EPU were used to acquire cryo-EM movies at Advanced Center for Electron Microscopy at Shanghai Institute of Materia Medica, Chinese Academy of Sciences. A total of 8,002 Movies were recorded with pixel size of 0.824 Å at a dose of 50 electron per Å^2^ for 36 frames. The defocus range of this dataset was −0.8 µm to −1.8 µm. For dimer complex, another 5,054 movies ware obtained with same parameters.

Cryo-EM images of the PTHrP-PTH1R-G_s_ complex were collected on a Titan Krios G4 at 300KV accelerating voltage equipped with Gatan K3 direct electron detector. A total of 7,266 movies were recorded with pixel size of 0.824 Å at a dose of 50 electron per Å^2^ for 36 frames. The defocus range of this dataset was −0.8 µm to −1.8 µm.

### Image processing

All dose-fractionated image stacks were subjected to beam-induced motion correction by RELION-4.0^23^. The defocus parameters were estimated by CTFFIND 4.1^24^. PTH-PTH1R-G_s_ complex and dimer complex, by auto-picking yielded 3,051,218 particles and 10,497,057 which were processed by reference-free 2D classification using Cryosparc^25^. With initial model, after several rounds of 3D classification using RELION, 407,467 particles and 55,858 particles were used to further refinement and polishing, yielding two reconstructions with global resolution of 2.62 Å and 3.72 Å respectively, and subsequently post-processed by DeepEMhancer^26^.

For the PTHrP-PTH1R-G_s_ complex, movies were aligned with RELION-4.0^23^. Initial contrast transfer function (CTF) fitting was performed with CTFFIND4.1^24^ from Cryosparc^25^. Auto-picking and two-dimensional (2D) Classification were processed using Cryosparc, producing 7,700,728 particles for further processing. With initial model, two rounds of 3D classifications were carried out. A further one round of 3D classification was conducted with a mask on the receptor, in which 436,359 particles were subjected to 3D auto-refinement and polishing. A map with an indicated global resolution of 3.25 Å at a Fourier shell correlation (FSC) of 0.143 was generated from the final 3D refinement, and subsequently post-processed by DeepEMhancer^26^.

### Model building and refinement

The cryo-EM structure of the LA-PTH1R-G_s_-Nb35 complex (PDB code 6NBF) and two ECD crystal structures (PDB: 3C4M and PDB: 3H3G) were used as the start for model building and refinement against the electron microscopy map. The model was docked into the electron microscopy density map using Chimera^27^, followed by iterative manual adjustment and rebuilding in COOT^28^. Real space and Rosetta refinements were performed using Phenix^29^. The model statistics were validated using MolProbity^30^. Fitting of the refined model to the final map was analyzed using model-versus-map FSC. To monitor the potential over-fitting in model building, FSC_work_ and FSC_free_ were determined by refining ‘shaken’ models against unfiltered half-map-1 and calculating the FSC of the refined models against unfiltered half-map-1 and half-map-2. The final refinement statistics are provided in Supplementary Table 2. Structural figures were prepared in Chimera and PyMOL (https://pymol.org/2/).

### Molecular dynamics simulations

The cryo-EM structures of PTH34-PTH1R-G_s_ complex and PTHrP-PTH1R-G_s_ complex were used for the construction of MD simulation systems. Missing ICL3 and short part before TM1 were supplied according to the loop builder program in Molecular Operation Environment. Missing N-terminal and ECL1 structures were constructed according to the AlphaFold2 models^31^. The mutation G188A was modified back and the G proteins were removed. Using CHARMM-GUI, the models were inserted in POPC (palmitoyl-2-oleoyl-sn-glycerol-3-phosphocholine) lipids^32,33^. The length of X and Y were both 130 Å. TIP3P waters and 0.15 M NaCl under a periodic boundary condition were also added to systems. The force fields for proteins and lipids were FF19SB and lipid17, respectively^34-36^. We used hydrogen mass repartitioning during system construction.

After preparation, the systems first encountered a minimization process of 5,000 steepest descent cycles with a constraint on backbone atom, sidechain atom, and lipid coordinates. Then, the constraints were generally decreased in the separated 6 steps of the equilibration process provided by CHARMM-GUI. The first three of them had a timestep of 1 fs and heated the system to 303.15 K, 1 atm in 375 ps. The last three of them had 2 fs timestep and decreased constrain in the coming 1.5 ns. We then performed the independent 400 ns × 6 MD simulations on Amber 20. During simulations, the Langevin thermostat and Berendsen barostat were used for temperature (300 K) and pressure (1 atm) control. Long-range electrostatic interactions were treated by the Particle mesh Ewald and a 10 Å cutoff was employed for short-range interactions. In the following analysis, tICA was calculated in MSMbuilder^37^ and the representative structures were extracted by CPPTRAJ^38^. MMGBSA were calculated by Maestro, Schrödinger.

### cAMP accumulation assay

PTH and PTHrP stimulated cAMP accumulation was measured by a LANCE Ultra cAMP kit (PerkinElmer). After 24 h culture, the transfected cells were seeded into 384-well microtiter plates at a density of 3,000 cells per well in HBSS supplemented with 5 mM HEPES, 0.1% (w/v) BSA or 0.1% (w/v) casein and 0.5 mM 3-isobutyl-1-methylxanthine. The cells were stimulated with different concentrations of peptide agonists for 40 min at RT. Eu-cAMP tracer and ULight™-anti-cAMP were then diluted by cAMP detection buffer and added to the plates separately to terminate the reaction. Plates were incubated at RT for 1 h and the fluorescence intensity measured at 620 nm and 665 nm by an EnVision multilabel plate reader (PerkinElmer).

### Statistical analysis

All functional data were displayed as means ± standard error of the mean (S.E.M.). Statistical analysis was performed using GraphPad Prism 8.0 (GraphPad Software). Experimental data were evaluated with a three-parameter logistic equation. The significance was determined with either two-tailed Student’s t-test or one-way ANOVA. *P* < 0.05 was considered statistically significant.

### Data availability

Cryo-EM maps have been deposited in the Electron Microscopy Data Bank under accession codes: EMD-XXX (PTH-bound PTH1R-G_s_ complex), EMD-XXX (PTH-bound PTH1R-G_s_ complex). The atomic coordinates have been deposited in the Protein Data Bank under accession codes: XXX (PTH-bound PTH1R receptor) and XXX (PTHrP-bound PTH1R receptor).

## RESULTS

### Structural determination

Stable PTH1R-G_s_ protein complex can be assembled with LA-PTH, but not with PTH and PTHrP. To overcome this technical obstacle, we employed the NanoBiT tethering strategy to stabilize the PTH1R-G_s_ protein complex assembly with PTH and PTHrP (Supplement Fig.1). The PTH1R was co-expressed with Gα_s_, Gβ1, Gγ2 and GST-Ric-8B. Nanobody 35 (Nb35) was added into membranes to stabilizing the receptor-G protein complex according to the method previously described^2^ (Supplement Fig. 1).

The complex structures of the G_s_-coupled PTH1R bound to PTH and PTHrP were determined by cryo-EM to the resolutions of 2.6 Å and 3.2 Å, respectively (Fig. 1 and Supplementary Fig. 2-3, Supplementary Table 1). In addition, we applied extensive particle classifications that yielded a distinct subclass of a dimeric form of the PTH-PTH1R-G_s_ complexes, which was determined to a resolution of 3.7 Å (Supplementary Fig. 4). Because this dimeric form is mediated through no-physiological Nb35 and G_s_ interface, no further discussion is presented in the paper. Both endogenous peptide-PTH1R-G_s_ complexes are similar to other GPCR-G protein complex structures, the α-helical domain (AHD) of G_s_ is flexible and invisible. Apart from extracellular domain (ECD) of receptor, the majority of the amino acid side chains of receptor and heterotrimeric G_s_ protein were well resolved in the final models, which are refined against the EM density map with excellent geometry. Both peptides, PTH (1-34) and PTHrP (1-36) were clearly identified, thus providing reliable models for the mechanistic explanation of pharmacological actions of teriparatide and abaloparatide at PTH1R (Supplement Fig. 5). In addition, like LA-PTH-PTH1R-G_s_ complex, an ordered annular lipid band wrapping around the periphery of the receptor transmembrane domain (TMD) is visible in the cryo-EM map, among which several lipid molecules were built (Fig. 1a-e). These structural lipids possibly stabilize the PTH1R in its active state.

**Fig. 1.**
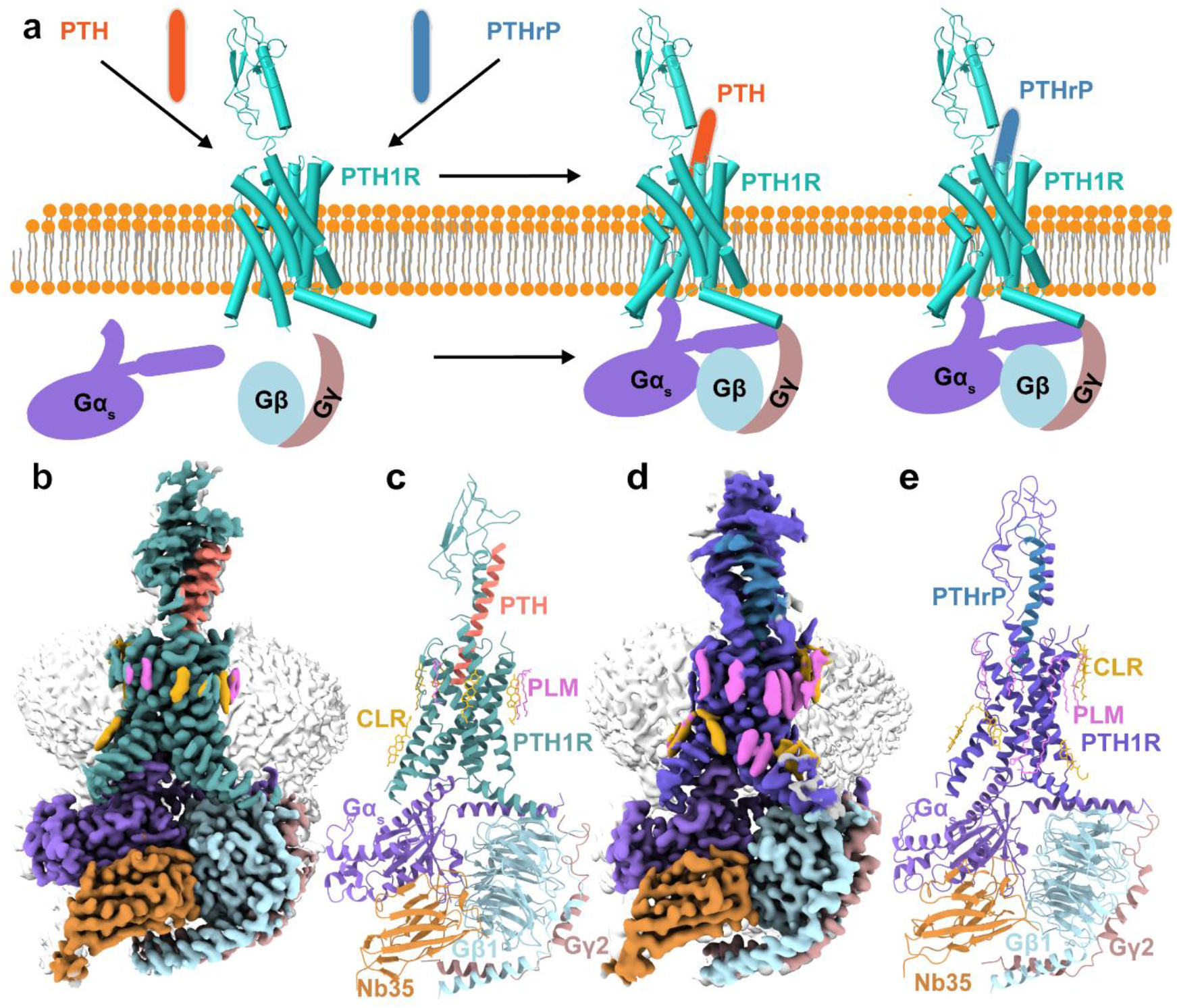
The overall cryo-EM structures of G_s_-coupled PTH1R bound to PTH and PTHrP. **(a)** Schematic diagram of two endogenous hormones, PTH (salmon) and PTHrP (steel blue) are recognized by PTH1R (light sea green) and exert their effects on bone by activating PTH1R, primarily via G_s_-cAMP signaling. G_s_ is colored in medium purple, Gβ in light blue and Gγ in rosy brown. **(b)** Orthogonal views of the density map and **(c)** model for PTH-PTH1R-G_s_ complex. The colored cryo-EM density map is shown at the 0.0183 threshold and light gray surface indicates a micelle diameter of 10 nm. Cadet blue, PTH1R; salmon, PTH; medium purple, G_s_; light blue, Gβ; rosy brown, Gγ; peru, Nb35; goldenrod, CLR; orchid, PLM. **(d)** Orthogonal views of the density map and **(e)** model for PTHrP-PTH1R-G_s_ complex. The colored cryo-EM density map is shown at the 0.136 threshold and light gray surface indicates a micelle diameter of 11 nm. Slate blue, PTH1R; steel blue, PTHrP; medium purple, G_s_; light blue, Gβ; rosy brown, Gγ; peru, Nb35; goldenrod, CLR; orchid, PLM.

### Overall structures of G_s_-coupled PTH1R bound to PTH and PTHrP

Both PTH-PTH1R-G_s_ and PTHrP-PTH1R-G_s_ complex structures closely resemble that of the LA-PTH-PTH1R-G_s_ complex with the root mean square deviation (RMSD) value of 0.5 Å for whole complexes, 0.7 Å for G_s_ alone, 0.5 Å for receptor alone (Fig. 2). Notable conformational differences were observed in the position and orientation of the C-terminal fragment of peptide and the ECD of receptor. The C-terminal fragment of PTH and PTHrP in complexes are shifted away TM7 by ∼9.6 Å and 2.6 Å (measured at Cα of H^32P^), respectively, relative to their positions in the LA-PTH-bound structure (PDB: 6NBF) ^2^. The receptor ECD-α1 helix in the PTH- and PTHrP-bound structures are shifted away TM7 by ∼10 Å and 7.1 Å (measured at Cα of K50^ECD^), respectively, relative to their positions in the LA-PTH-bound structure (Fig. 2a-c). These conformational changes allow the C-terminal fragment of hormone bind specificity to PTH1R, indicating PTH1R-associated peptide recognition specificity. Except ECD-α1 helix in PTH-PTH1R cryo-EM structure is shifted away PTH by ∼2 Å, relative to X-ray structure (measured at Cα of K50^ECD^), the overall conformation of the PTH-receptor ECD and PTHrP-receptor ECD in PTH-PTH1R cryo-EM structures resemble the previously published crystal structures of PTH-ECD and PTHrP-ECD complexes (PDB: 3C4M and 3H3G) ^9,15^, respectively. The structural basis for the binding specificity of PTH and PTHrP to PTH1R is relatively consistent with the crystal structures (Fig. 2d).

**Fig. 2.**
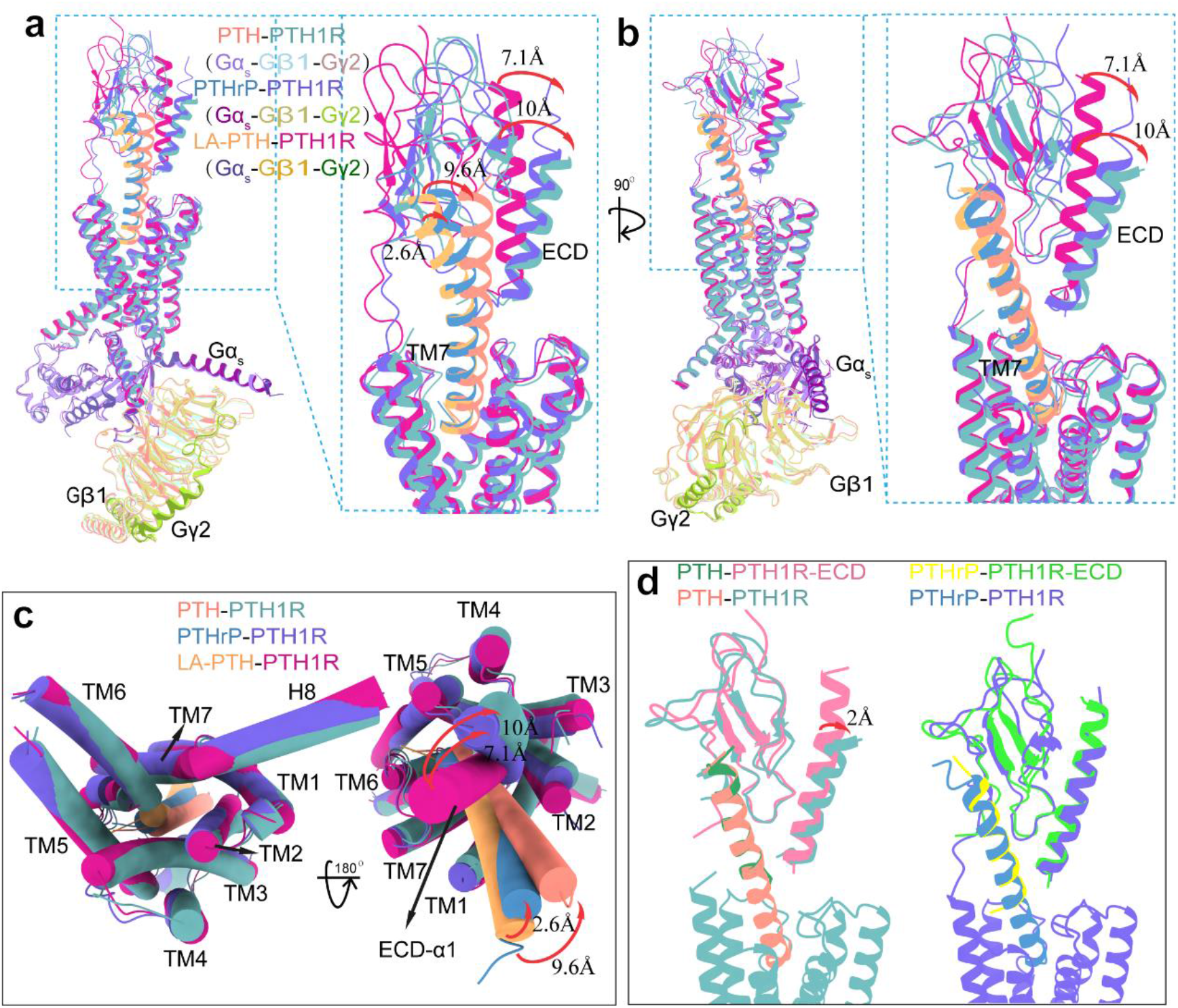
Structure comparisons of PTH1R-G_s_ complexes by different ligands. **(a-c)** Structural comparison of PTH-PTH1R-G_s_ (salmon, PTH; cadet blue, PTH1R; medium purple, G_s_; light blue, Gβ; rosy brown, Gγ), PTHrP-PTH1R-G_s_ (steel blue, PTHrP; slate blue, PTH1R; purple, G_s_; dark khaki, Gβ; yellow green, Gγ) and LA-PTH-PTH1R-G_s_ (PDB: 6NBF, sandy brown, LA-PTH; medium violet red, PTH1R; dark slate blue, G_s_; goldenrod, Gβ; dark green, Gγ) complexes. Notable conformational changes observed in the receptor ECD and the ligand **(a-b)** in the side view and **(c)** cytoplasmic and extracellular view. **(d)** Comparison of the PTH bound ECD in PTH-PTH1R-G_s_ EM structure with PTH-bound ECD crystal structure (PDB: 3C4M) and PTHrP bound ECD in PTHrP-PTH1R-G_s_ EM structure with PTHrP-bound ECD crystal structure (PDB: 3H3G), yellow, PTHrP; lime green, ECD; sea green, PTH; pale violet red, ECD.

### Molecular basis for recognition of PTH and PTHrP by PTH1R

Both PTH-PTH1R-G_s_ and PTHrP-PTH1R-G_s_ complexes exhibit a similar and unique peptide-receptor binding interface, which is consistent with the “two-domain” model of the peptide-PTH1R interaction mechanism of hormone recognition common to class B GPCRs (Fig. 3). In these complexes, PTH and PTHrP adopt an amphipathic helix with their N terminus dipping into the receptor TMD, while the C terminus is closely interaction with the ECD. The N-terminal fragment of PTH (residues 1-14, PTH^N^) and PTHrP (residues 1-15, PTHrP^N^) bind to the TMD and activate the receptor. The PTH^N^ and PTHrP^N^ adopt four helical turns and insert deeply into the TMD pocket. PTH^N^ and PTHrP^N^ are directly surrounded by TM1, TM2, TM3, TM5, TM6 and TM7 and form extensive interactions with the TMD core. PTH^N^ and PTHrP^N^ can also interact with ECL2 and ECL3 of receptor. The C-terminal fragment of PTH (residues 15-34, PTH^C^) and PTHrP (residues 16-34 PTHrP^C^) bind to the ECD of PTH1R to confer high affinity and specificity to the receptor (Fig. 3a, d).

**Fig. 3.**
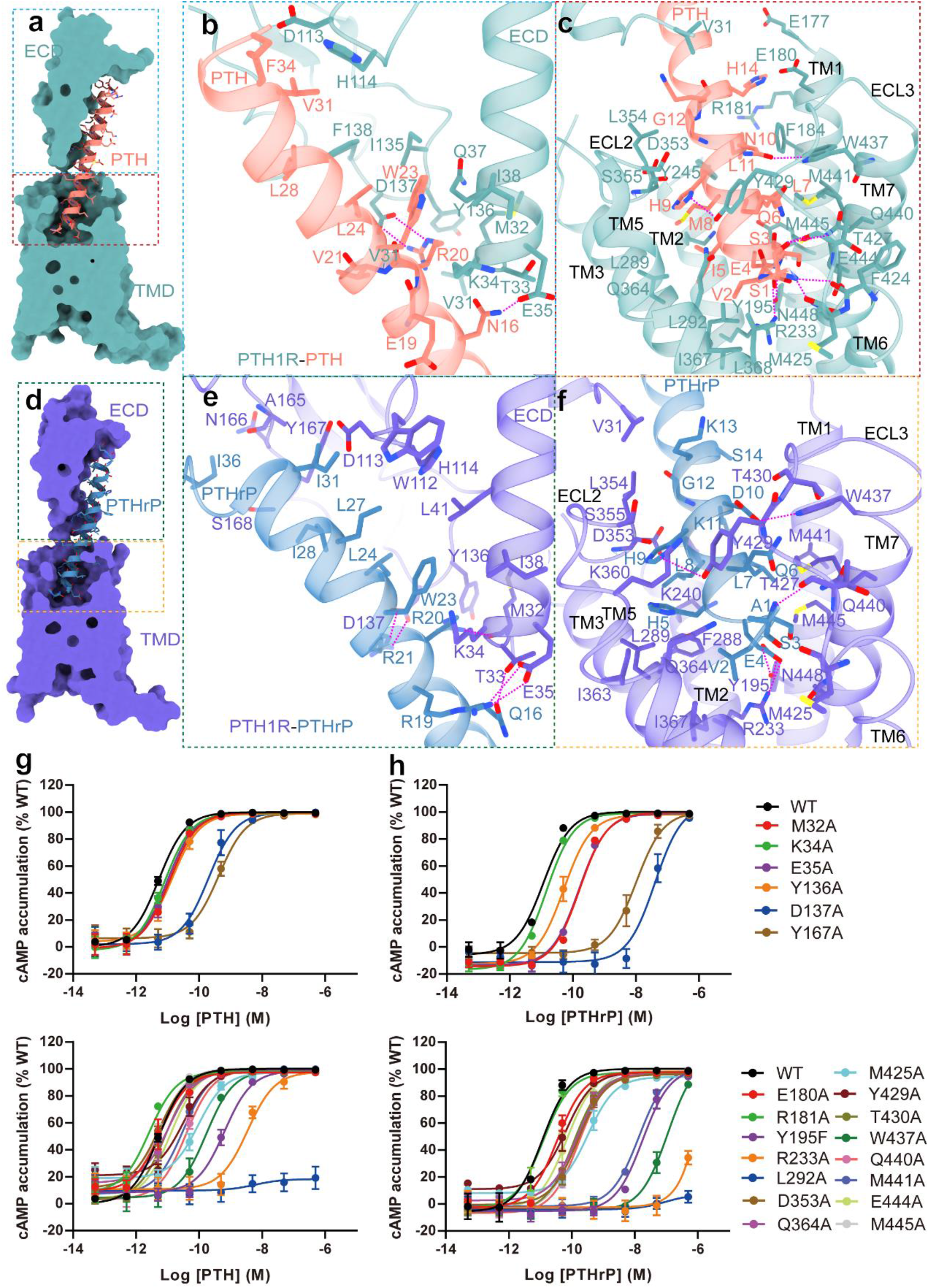
Molecular recognition of PTH1-34 and PTHrP1-36 by PTH1R. **(a)** An overall PTH-binding pocket of PTH1R. Cutaway view showing that PTH^Nα^ penetrates into a pocket formed by TMD, while its C-terminal half is recognized by the ECD. **(b-c)** Detailed interactions of PTH^C^ with the PTH1R ECD and interactions of PTH^Nα^ with the PTH1R TMD pocket, salmon, PTH; cadet blue, PTH1R. The polar contacts are shown as purple dotted lines. **(d)** An overall PTHrP-binding pocket of PTH1R. Cutaway view showing that PTHrP^Nα^ penetrates into a pocket formed by TMD, while its C-terminal half is recognized by the ECD. **(e-f)** Detailed interactions of PTHrP^C^ with the PTH1R ECD and interactions of PTHrP^Nα^ with the PTH1R TMD pocket, steel blue, PTHrP; slate blue, PTH1R. The polar contacts are shown as purple dotted lines. **(g-j)** Effect of receptor mutation on PTH and PTHrP-induced cAMP accumulation. **(g-h)** Signaling profiles of PTH1R ECD mutants on PTH and PTHrP-induced cAMP accumulation. **(i-j)** Signaling profiles of PTH1R TMD, ECL2 and ECL3 mutants on PTH and PTHrP-induced cAMP accumulation. cAMP accumulation was measured in wild-type (WT) and single-point mutated PTH1R expressing in HEK293T cells, respectively. cAMP accumulation was normalized to the maximum response of the WT and dose-response curves were analyzed using a three-parameter logistic equation. Data were generated and graphed as means ± S.E.M. of three independent experiments (n = 3) performed in triplicate.

In line with a similar conformation observed between the cryo-EM structures and crystal structures of the PTH1R ECD bound PTH and PTHrP, the extensive interactions of the ECD with PTH^C^ and PTHrP^C^ in cryo-EM structure complexes are similar to those observed in the previously reported X-ray structures of peptide-ECD complexes (Fig. 3b, e). In the PTH-PTH1R-G_s_ complex, the hydrophobic residues V21, W23, L24, L28, V31 and F34 of the PTH, form an extensive hydrophobic interactions with the hydrophobic groove of the three-layer PTH1R-ECD fold (Fig. 3b)^15^. In the PTHrP-PTH1R-G_s_ complex, the conserved ECD fold also shows the hydrophobic contacted with F23, L24, L27, I28, I31 and I36 of PTHrP (Fig. 3e)^9^. In addition, N16^PTH^ and Q16^PTHrP^ form a hydrogen bond with E35^ECD^ and T33^ECD^, respectively. The conserved amino acid R20 in the PTH and PTH-related ligands also forms a hydrogen bond with the backbone carbonyls of M32^ECD^ of PTH1R and forms electrostatic interactions with D137^ECD^. These polar interactions display the binding affinity and specificity of PTH and PTHrP to PTH1R. This amino acid R20 is conserved in all three endogenous peptide hormones of PTH1R and PTH2R but distinct from other class B GPCRs, suggesting that R20 is critical for peptide hormones recognition and specificity of PTH and PTH-related ligands^15^. Except for these conserved interaction, these two peptide hormones also interact with PTH1R in a peptide-specific mode. E19^PTH^ forms electrostatic interaction with K34^ECD^, V21^PTH^ makes hydrophobic contact with D137^ECD^ and W23^PTH^ makes Pi interaction with K34^ECD^ and hydrophobic interaction with Q37^ECD^ and I38^ECD^. But different amino acids at cognate positions of PTHrP, R19^PTHrP^ and R21^PTHrP^ form electrostatic interaction with E35^ECD^, D137^ECD^, respectively and F23^PTHrP^ makes hydrophobic interaction with I38^ECD^ and L41^ECD^ of PTH1R (Fig. 3b, e). These interactions receive support from our mutagenesis studies by measuring cAMP responses. Alanine substitution of D137^ECD^ and Y167^ECD^ of PTH1R shows clearly a great reduction in the potency of both PTH and PTHrP-mediated G_s_ activation (Fig. 3g-h, Supplementary Table 2). In additional, both alanine substitution M32^ECD^ and E35^ECD^ also diminish the potency of PTHrP and slightly decreased the efficacy of PTH. These amino acid residues are important for determining the affinity and specificity of PTH and PTHrP to PTH1R.

The sequences of PTH^N^, PTHrP^N^ and other PTH-related ligands highly resemble. Although PTHrP^N^ can overlapped well with LA-PTH^N^, while PTH^N^ is different from their positions. Every one of the three N-terminal peptide helix inserts into the receptor TMD core by a analogical angle and orientation, thereby revealing a similar peptide hormones recognition and activation mechanism (Fig. 2c, f and supplementary Fig. 6). Both PTH^N^ and PTHrP^N^ run parallel to TM2, making extensive contacts with the residues in TM1, TM2, TM3, TM5, TM6, TM7 and ECL2 and ECL3 (Fig. 3c, f). Especially the first six residues of PTH ^N^ and PTHrP ^N^ play critical roles in the receptor activation. In contrast, PTH (7-34) and PTHrP (7-34) are potent antagonists of PTH receptor^1^. For PTH, A1 forms a hydrogen bond with T427^6.59b^ and makes hydrophobic contacts with Q364^5.40b^, M425^6.57b^ and Q440^7.38b^. For PTHrP, S1 makes more polar interactions with F424^6.56b^, M425^6.57b^, T427^6.59b^ and Q440^7.38b^. These observations provide support for our mutagenesis studies by measuring cAMP responses. Alanine substitutions of Q364^5.40b^, M425^6.57b^ and Q440^7.38b^ notably reduce the potency PTHrP-mediated G_s_ activation, and slightly effect on the potency of PTH in G_s_ activation. (Fig. 3i-g, Supplementary Table 2). The same amino acid V2 in two peptides makes extensive hydrophobic contacts with L292^3.40b^, Q364^5.40b^ and I367^5.43b^. Mutation of L292^3.40b^ remarkably reduce pEC_50_ and E_max_ values in the PTH- and PTHrP-induced receptor activity (Fig. 3i-g, Supplementary Table 2). Although S3 in the same position, they exhibit different binding interaction with receptor. S3^PTH^ makes hydrophobic contacts with Q440^7.38b^, M441^7.39b^ and M445^7.43b^. While S3^PTHrP^ forms polar interactions with E444^7.42b^, N448^7.46b^, and also makes hydrophobic interaction with M445^7.43b^. alanine substitutions of M445^7.43b^ indicates a great reduction in the potency of PTHrP-mediated G_s_ activation (Fig. 3j, Supplementary Table 2), but almost no effect on the receptor activity by PTH in G_s_ activation (Fig. 3i, Supplementary Table 2). E4 in both PTH and PTHrP, forms the stabilizing hydrogen bond interact and salt bridge with Y195^1.47b^ and R233^2.60b^, respectively. Both alanine mutations in Y195^1.47b^ and R233^2.60b^ remarkably reduce peptide-mediated PTH1R activation, indicating that E4 of the peptide is a key role for peptide-induced receptor activation (Fig. 3i-j, Supplementary Table 2). I5^PTH^ makes hydrophobic contacts with L289^3.37b^, L292^3.40b^ and Q364^5.40b^. While H5^PTHrP^ at the same position of PTHrP not only makes extensive hydrophobic interactions with L289^3.37b^, K360^5.36b^ and I363^5.39b^, but also forms an additional electrostatic interaction with Q364^5.40b^. Q6 of PTH and PTHrP forms highly conserved hydrophobic interactions with Y429^ECL3^, W437^7.35b^, Q440^7.38b^ and M441^7.39b^. All of these residues mutations, Y429^ECL3^A, W437^7.35b^A, Q440^7.38b^A and M441^7.39b^A reduce both PTH and PTHrP to activate PTH1R. In additional, N10^PTH^ is hydrogen-bonded with W437^7.35b^, the same to PTH position, D10^PTHrP^ also forms a hydrogen-bond with W437^7.35b^. W437^7.35b^A, markedly reduce both PTH and PTHrP to activate PTH1R, supporting the fact that N10^PTH^ and D10^PTHrP^ of the peptide are critical for two endogenous hormones-induced receptor activation (Fig. 3i-j, Supplementary Table 2). H9 in both PTH and PTHrP interacts with ECL2 and ECL3 of receptor, such as the hydrophobic interactions with D353^ECL2^, S355^ECL2^, and electrostatic interaction with Y429^ECL2^. D353^ECL2^ significantly decreases the PTHrP-induced PTH1R activation (Fig. 3j, Supplementary Table 2), but almost no effect on the receptor activity by PTH in G_s_ activation (Fig. 3i, Supplementary Table 2). In additional, L7^PTH/PTHrP^, M8^PTH^ or L8^PTHrP^, L11^PTH^ or K11^PTHrP^, G12^PTH/PTHrP^, K13^PTH/PTHrP^, H14^PTH^ or K14^PTHrP^ can form extensive hydrophobic interactions with the TMD core and ECL2. The above interactions of peptide^N^ with the TMD provide the major binding and activation energy for the formation of the signal complex. Therefore, the stability of the interface between peptide^N^ and the TMD is well supported by an extensive network of complementary polar and non-polar interactions between the bound PTH and PTH-related ligands and the TMD. The detailed interactions listed in Fig. 2c, f and supplementary Fig. 6. These interactions help to explain the rich and distinct pharmacology of PTH and PTHrP acting through the same receptor.

### Activation mechanism of PTH1R

Comparison of the PTH- and PTHrP-bound PTH1R-G_s_ complex structures with LA-PTH-bound PTH1R-G_s_ complex structure reveals a remarkably similarity in the G protein binding interface, indicating a common mechanism for G protein coupling specificity (Fig. 4). Gα_s_-α5 helix dipped into the cytoplasmic receptor cavity and forms important interactions with TM2, TM3, TM5, TM6, H8 and intracellular loop 3(ICL3) (Fig. 4c-d). In all three PTH1R-G_s_ complex structures, L393 in Gα_s_-α5 helix make polar interaction with S409^6.41b^. Both D381 and Q384 of Gα_s_-α5 helix form an electrostatic interaction with K388^5.64b^. R385 of Gα_s_-α5 also makes an additional electrostatic interaction with E391^ICL3^ of PTH1R in PTH-bound PTH1R-G_s_ complex and PTHrP-bound PTH1R-G_s_ complex. E392 forms polar interaction with the backbone amine of N363^8.47b^ and G364^8.48b^. The bulky residue Y391 at the Gα_s_-α5 helix C terminus binds to a sub-pocket formed by R219^2.46b^, H233^2.50b^, Y305^3.53b^ and L306^3.54b^ in PTH1R. The hydrophobic residues L388, Y391, L393 and L394 of Gα_s_-α5 helix form extensive hydrophobic interaction with TMD helixes (Fig. 4c-d). F314^ICL2^ in PTH- and PTHrP-bound PTH1R-G_s_ complexes insert into a cavity formed by αN-β1 junction and α5 of Gα_s_ and make hydrophobic interactions for stabilizing the interface, but they are different from LA-PTH-bound PTH1R-G_s_ complex, which F314^ICL2^ not directly interact with the hinge region of the Gα_s_ protein (Fig. 4e). Both in PTH- and PTHrP-bound PTH1R-G_s_ complex, the electrostatic interactions presented between R214^ICL1^ and D312 of Gβ1, and in PTH- and LA-PTH-bound PTH1R-G_s_ complex, hydrogen bonds between R476^8.60b^ and the backbone oxygen of G310 of Gβ1, while in PTHrP-bound PTH1R-G_s_ complex, hydrogen bonds between R472^8.56b^ and the backbone oxygen of G310 of Gβ1. Additional contacts between R213^ICL1^ and R52 of Gβ1 in PTH-bound PTH1R-G_s_ complex and in LA-PTH-bound PTH1R-G_s_ complex, all of these interactions may stabilize the interface of helix 8 and Gβ1 (Fig. 4f).

**Fig. 4.**
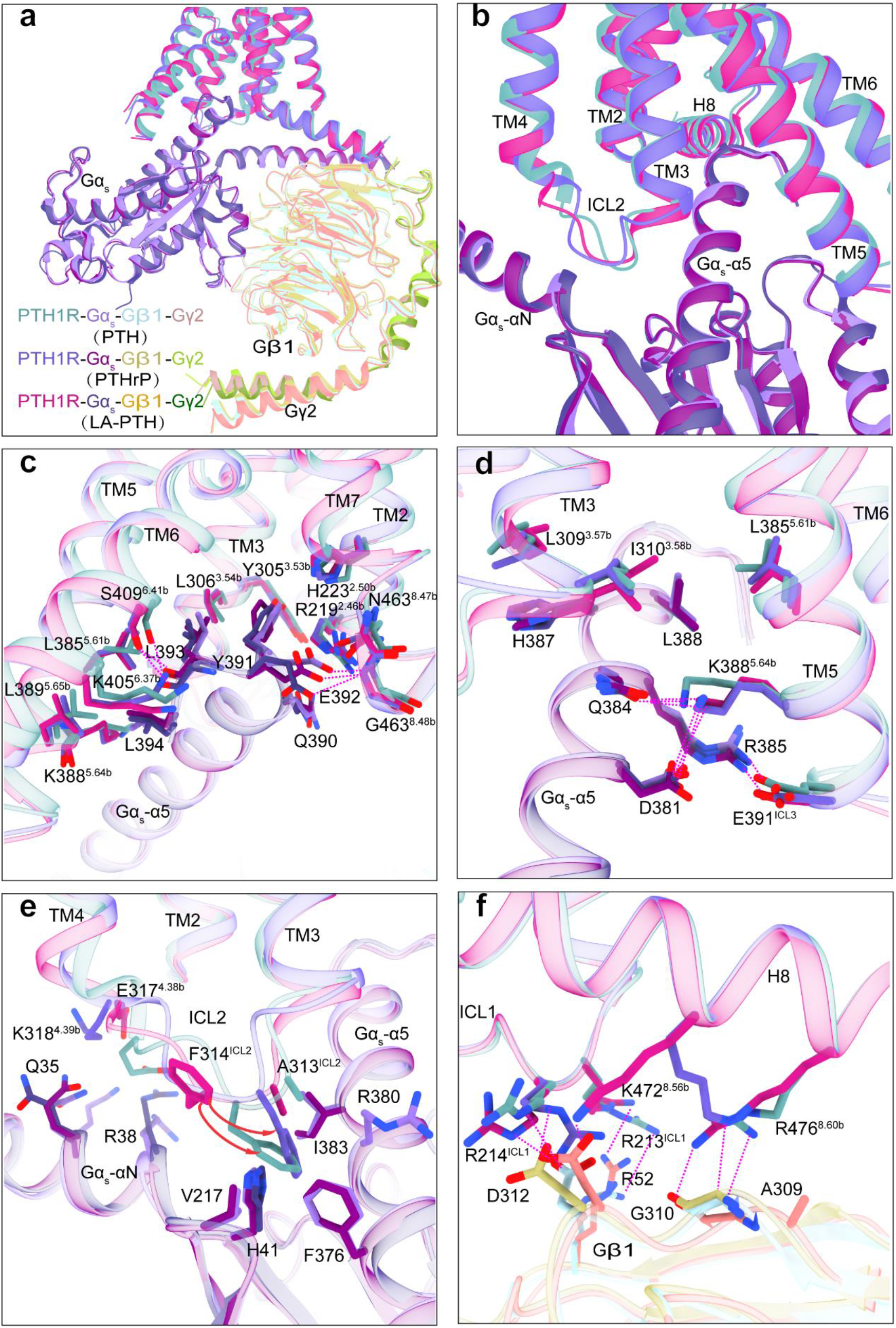
G protein coupling of multi-agonist-bound PTH1R. **(a)** Comparison of G protein coupling among PTH, PTHrP and LA-PTH bound PTH1R. The receptors and G proteins are colored as the labels. **(b)** The Gα_s_-α5 helix of the Gα_s_ inserts into anintracellular cavity of receptor’s TMD. **(c-f)** Comparison of the PTH1R-G_s_ interface, including **(c)** the Gα_s_-α5 C terminus-TMD interface, **(d)** Gα_s_-α5-TM3/TM5 interface, **(e)** ICL2-Gα_s_-αN and Gα_s_-α5 helix interface, **(f)** and helix 8-Gβ1 interface. cadet blue, PTH1R; medium purple, G_s_; light blue, Gβ; rosy brown, Gγ in PTH-PTH1R-G_s_ complex, slate blue, PTH1R; purple, G_s_; dark khaki, Gβ; yellow green, Gγ in PTHrP-PTH1R-G_s_ complex and medium violet red, PTH1R; dark slate blue, G_s_; goldenrod, Gβ; dark green, Gγ in LA-PTH-PTH1R-G_s_ complex The polar contacts are shown as purple dotted lines.

### Conformational changes during PTH1R activation

PTH1R has two different conformation states, the G-protein free state R^0^ and the G protein coupled state RG^10^. During peptide-bound PTH1R activation, two PTH1R conformations undergo different rearrangement of the network residues to facilitate the ligand binding and signal transduction^14^. Based on structural alignment of PTH1R: agonist complex in the absence of G protein, with three G protein-bound PTH1R (RG): agonist complexes shows different conformational changes, involving the ECD and the TMD of the receptor (Fig. 5a-c). The receptor ECD-α1 in the PTH-bound and PTHrP bound PTH1R RG structures are shifted away TM7 by ∼4.4 Å and 3.5 Å (measured at Cα of K50^ECD^), respectively, relative to their positions in the ePTH-bound PTH1R structure. The main conformational changes happens in TMD (Fig. 5b-c). The class B GPCRs conserved PxxG motif (P415^6.48b^, L416^6.48b^, F417^6.49b^ and G418^6.50b^) in three agonist-G-protein-bound PTH1R complexes caused a notable conformational rearrangement and make TM6 kink outward ∼90° in the middle (Fig. 5d). The TM6 sharp kink creating an intracellular binding pocket for the C terminus of Gα, thereby enabling intracellular G protein activation and signal transduction. Meanwhile, the kink of TM6 together with other conformational changes caused three class B GPCRs conserved polar interaction networks rearrangement, including HETY network (H223^2.50b^, E302^3.50b^, T410^6.42b^, and Y457^7.57b^), the II-VI-VII-VIII network (R219^2.46b^, K405^6.37b^, N463^7.61b^ and E465^8.49b^) and central polar network (R233^2.60b^, N295^3.43b^, H420^6.52b^ and Q451^7.49b^). However, this structural hallmark of class B GPCRs activation was not observed in the ePTH-PTH1R complex structure and CLR-CGRPR-RAMP1 complex structure, which are in the absence of G protein. Their intracellular part of the receptors still displays the hallmarks of an inactive conformation (Fig. 5b and supplementary Fig. 7), which are called intermediate states on the transition of the receptor toward the G protein-bound state^13,14,16^. The intermediate states of class B GPCRs and the R^0^ conformation of the PTH1R resemble in agonist binding and in the absence of G protein. The ability of PTH and PTHrP to bind to RG is similar, but PTH has a greater capacity to bind to R^0^ state^1,10^. Such conformational selectivity mediate their distinct biological actions^10^. From molecular dynamics (MD) simulations, the lower R^0^ binding ability of PTHrP was attributed to its higher binding free energy upon specific states (Supplementary Fig. 8). Although the capacity of PTH and PTHrP to bind to R^0^ is different, these peptides can bind to the G protein-coupled PTH1R complex (RG) with a highly similar G protein binding interface, suggesting a common mechanism for G_s_ engagement.

**Fig. 5.**
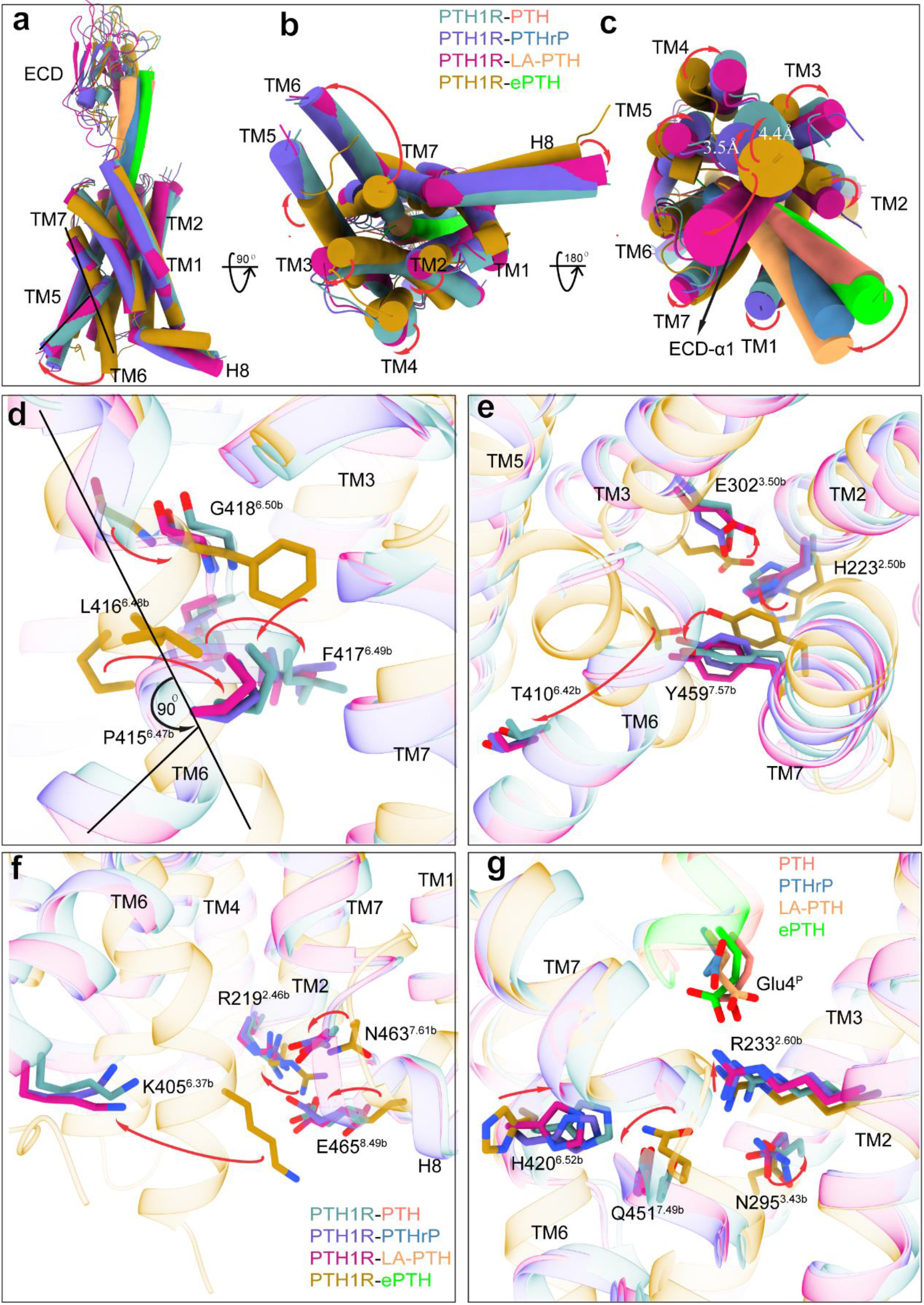
Structures of the active PTH1R in G protein-bound state and comparison with structure of the active PTH1R in the absence of G protein. **(a-c)** The structural alignment of ePTH-bound PTH1R (PDB: 6FJ3, lime, ePTH; dark goldenrod, PTH1R) with PTH-bound PTH1R (salmon, PTH; cadet blue, PTH1R), PTHrP-bound PTH1R (steel blue, PTHrP; slate blue, PTH1R) and LA-PTH-bound PTH1R (sandy brown, LA-PTH; medium violet red, PTH1R), which are in G_s_ bound state, showing different conformational changes of the receptor between PTH1R in G_s_ bound and no G_s_ bound states **(a)** in the side view, **(b)** extracellular view and **(c)** intracellular view, the red arrows indicate conformational changes. **(d)** The structural alignment of PTH1R in the absence of G protein and in G protein-bound state, showing the outward bending of the intracellular portion of TM6 of activated PTH1R in G_s_-bound state, which results in a kink at the PxxG motif in TM6 and a ∼90° angle in the middle of TM6 of the activated PTH1R in G protein-bound state. The TM6 kink in the active PTH1R structure is indicated by a black line. The conserved PxxG motif in TM6 are shown in stick representation. **(e)** Conformational changes during receptor activation in the absence of G protein and in G protein-bound state in HETY network, **(f)** in the II-VI-VII-VIII network, **(g)** and in central polar network. The residues are shown in stick representation and conformational changes are indicated by red arrows.

## DISCUSSION

The cryo-EM structures of PTH-PTH1R-G_s_ and PTHrP-PTH1R-G_s_ complexes further reveal the structural basis for recognition of PTH and PTHrP by PTH1R and the receptor activation and G protein-coupling mechanisms, providing an opportunity to use these structures as modules for designing new PTH replacement therapies in the future. Two synthetic analogs of PTH and PTHrP, teriparatide and abaloparatide were approved drugs by FDA and used in clinic, but as conventional therapy for osteoporosis by daily injections with high costs, probably not suitable for treating such a widespread disease. Therefore, PTH1R-specific targeted orally agonists for osteoporosis are urgently developed. To our knowledge, PTH and PTHrP differ in selectivity for two pharmacologically distinguishable high-affinity PTH1R conformations, R^0^ and RG^1^. Such R^0^-selective analogs, including PTH, M-PTH (1-28) and M-PTH (1-34) display high affinity binding to R^0^ as well as notably mediate prolonged cAMP signaling responses and are now being investigated as potential new drugs for treating hypoparathyroidism^17^. While another PTH analogs, such as PTHrP (1-36) and M-PTH (1-14) exhibit a high-affinity to bind to RG conformation, and produces a shorter-lived and more pulsatile action at the receptor, might prove to be more efficacious in this class of patients^1,12^. Mutational studies of the ligand-receptor binding sites indicate that the binding specificity of PTH and PTHrP to PTH1R. According to our structures, detailed patterns of teriparatide and abaloparatide-receptor interactions can be identified and it can explain the binding specificity of PTH and PTHrP to PTH1R. Thus, a detailed understanding of PTH and PTHrP binding and activation of PTH1R provide a rational template to development of optimized therapeutics targeting PTH1R diseases.

## ACKNOWLEDGMENTS

The cryo-EM data were collected at Advanced Center for Electron Microscopy at Shanghai Institute of Materia Medica, Chinese Academy of Sciences. We are grateful to W.H. and K.W. for collecting the cryo-EM data. This work was supported by National Natural Science Foundation of China (32071203 to L.H.Z.; 82073904 to M.W.W. and 81973373 to D.H.Y.), the National Key R&D Program of China (2019YFA0904200), the Young Innovator Association of CAS (2018325 to L.H.Z.) and SA-SIBS Scholarship Program to L.H.Z. and D.H.Y.; Ministry of Science and Technology (China) grants (2018YFA0507002 to H.E.X. and 2018YFA0507000 to M.W.W.), the Shanghai Municipal Science and Technology Major Project (2019SHZDZX02 to H.E.X.;18ZR1447800 to L.H.Z and 21JC1401600 to D.H.Y.), the CAS Strategic Priority Research Program (XDB08020303 to H.E.X.).

## AUTHOR CONTRIBUTIONS

L.H.Z. designed the expression constructs, purified the complexes, prepared the final samples for negative stain and data collection toward the structures, participated in model building and performed structure and function data analysis, prepared figures and wrote the manuscript; L.H.Z. prepared the cryo-EM grids and Q.N.Y. collected cryo-EM images and performed map calculations, built and refined the structure models; Y.W.X. participated in model refinement; X.H.H conducted MD simulations; A.T.D., C.W.C., C.Z. and Y.Z. performed signaling experiments under the supervision of D.H.Y. and M.W.W.; L.H.Z. and H.E.X. conceived the project, wrote the manuscript.

## REFERENCES

1 Gardella, T. J. & Vilardaga, J. P. International Union of Basic and Clinical Pharmacology. XCIII. The parathyroid hormone receptors--family B G protein-coupled receptors. Pharmacol Rev 67, 310–337, doi:10.1124/pr.114.009464 (2015).

2 Zhao, L. H. et al. Structure and dynamics of the active human parathyroid hormone receptor-1. Science 364, 148–153, doi:10.1126/science.aav7942 (2019).

3 Nishimura, Y. et al. Lead Optimization and Avoidance of Reactive Metabolite Leading to PCO371, a Potent, Selective, and Orally Available Human Parathyroid Hormone Receptor 1 (hPTHR1) Agonist. J Med Chem 63, 5089–5099, doi:10.1021/acs.jmedchem.9b01743 (2020).

4 Kir, S. et al. Tumour-derived PTH-related protein triggers adipose tissue browning and cancer cachexia. Nature 513, 100–104, doi:10.1038/nature13528 (2014).

5 Wider, J., Undyala, V. V. R., Lanske, B., Datta, N. S. & Przyklenk, K. Parathyroid Hormone-Related Peptide and Its Analog, Abaloparatide, Attenuate Lethal Myocardial Ischemia-Reperfusion Injury. J Clin Med 11, doi:10.3390/jcm11092273 (2022).

6 Chew, C. K. & Clarke, B. L. Abaloparatide: Recombinant human PTHrP (1-34) anabolic therapy for osteoporosis. Maturitas 97, 53–60, doi:10.1016/j.maturitas.2016.12.003 (2017).

7 Srivastava, M. & Deal, C. Osteoporosis in elderly: prevention and treatment. Clin Geriatr Med 18, 529–555, doi:10.1016/s0749-0690(02)00022-8 (2002).

8 Bilezikian, J. P., Rubin, M. R. & Finkelstein, J. S. Parathyroid hormone as an anabolic therapy for women and men. J Endocrinol Invest 28, 41–49 (2005).

9 Pioszak, A. A., Parker, N. R., Gardella, T. J. & Xu, H. E. Structural basis for parathyroid hormone-related protein binding to the parathyroid hormone receptor and design of conformation-selective peptides. The Journal of biological chemistry 284, 28382–28391, doi:10.1074/jbc.M109.022905 (2009).

10 Dean, T., Vilardaga, J. P., Potts, J. T., Jr. & Gardella, T. J. Altered selectivity of parathyroid hormone (PTH) and PTH-related protein (PTHrP) for distinct conformations of the PTH/PTHrP receptor. Mol Endocrinol 22, 156–166, doi:10.1210/me.2007-0274 (2008).

11 Hattersley, G., Dean, T., Corbin, B. A., Bahar, H. & Gardella, T. J. Binding Selectivity of Abaloparatide for PTH-Type-1-Receptor Conformations and Effects on Downstream Signaling. Endocrinology 157, 141–149, doi:10.1210/en.2015-1726 (2016).

12 Okazaki, M. et al. Prolonged signaling at the parathyroid hormone receptor by peptide ligands targeted to a specific receptor conformation. Proc Natl Acad Sci U S A 105, 16525–16530, doi:10.1073/pnas.0808750105 (2008).

13 Ehrenmann, J. et al. High-resolution crystal structure of parathyroid hormone 1 receptor in complex with a peptide agonist. Nat Struct Mol Biol 25, 1086–1092, doi:10.1038/s41594-018-0151-4 (2018).

14 Ehrenmann, J., Schoppe, J., Klenk, C. & Pluckthun, A. New views into class B GPCRs from the crystal structure of PTH1R. Febs Journal 286, 4852–4860, doi:10.1111/febs.15115 (2019).

15 Pioszak, A. A. & Xu, H. E. Molecular recognition of parathyroid hormone by its G protein-coupled receptor. Proc Natl Acad Sci U S A 105, 5034–5039, doi:10.1073/pnas.0801027105 (2008).

16 Josephs, T. M. et al. Structure and dynamics of the CGRP receptor in apo and peptide-bound forms. Science 372, doi:10.1126/science.abf7258 (2021).

17 Mannstadt, M. et al. Efficacy and safety of recombinant human parathyroid hormone (1-84) in hypoparathyroidism (REPLACE): a double-blind, placebo-controlled, randomised, phase 3 study. Lancet Diabetes Endocrinol 1, 275–283, doi:10.1016/S2213-8587(13)70106-2 (2013).

18 Maeda, S. et al. Development of an antibody fragment that stabilizes GPCR/G-protein complexes. Nat Commun 9, 3712, doi:10.1038/s41467-018-06002-w (2018).

19 Chan, P. et al. Purification of heterotrimeric G protein alpha subunits by GST-Ric-8 association: primary characterization of purified G alpha(olf). The Journal of biological chemistry 286, 2625–2635, doi:10.1074/jbc.M110.178897 (2011).

20 Dixon, A. S. et al. NanoLuc Complementation Reporter Optimized for Accurate Measurement of Protein Interactions in Cells. ACS Chem Biol 11, 400–408, doi:10.1021/acschembio.5b00753 (2016).

21 Zhao, L.-H. et al. Structure and dynamics of the active human parathyroid hormone receptor-1. Science 364, 148–153, doi:10.1126/science.aav7942 (2019).

22 Ma, S. et al. Molecular Basis for Hormone Recognition and Activation of Corticotropin-Releasing Factor Receptors. Mol Cell 77, 669–680 e664, doi:10.1016/j.molcel.2020.01.013 (2020).

23 Zivanov, J., Nakane, T. & Scheres, S. H. W. Estimation of high-order aberrations and anisotropic magnification from cryo-EM data sets in RELION-3.1. Iucrj 7, 253–267, doi:10.1107/S2052252520000081 (2020).

24 Rohou, A. & Grigorieff, N. CTFFIND4: Fast and accurate defocus estimation from electron micrographs. J Struct Biol 192, 216–221, doi:10.1016/j.jsb.2015.08.008 (2015).

25 Punjani, A., Rubinstein, J. L., Fleet, D. J. & Brubaker, M. A. cryoSPARC: algorithms for rapid unsupervised cryo-EM structure determination. Nat Methods 14, 290–296, doi:10.1038/nmeth.4169 (2017).

26 Sanchez-Garcia, R. et al. DeepEMhancer: a deep learning solution for cryo-EM volume post-processing. Commun Biol 4, 874, doi:10.1038/s42003-021-02399-1 (2021).

27 Pettersen, E. F. et al. UCSF Chimera--a visualization system for exploratory research and analysis. J Comput Chem 25, 1605–1612, doi:10.1002/jcc.20084 (2004).

28 Emsley, P. & Cowtan, K. Coot: model-building tools for molecular graphics. Acta Crystallogr D Biol Crystallogr 60, 2126–2132, doi:10.1107/S0907444904019158 (2004).

29 Adams, P. D. et al. PHENIX: a comprehensive Python-based system for macromolecular structure solution. Acta Crystallogr D Biol Crystallogr 66, 213–221, doi:10.1107/S0907444909052925 (2010).

30 Chen, V. B. et al. MolProbity: all-atom structure validation for macromolecular crystallography. Acta Crystallogr D 66, 12–21, doi:10.1107/S0907444909042073 (2010).

31 Jumper, J. et al. Highly accurate protein structure prediction with AlphaFold. Nature 596, 583–589, doi:10.1038/s41586-021-03819-2 (2021).

32 Jo, S. et al. CHARMM-GUI 10 years for biomolecular modeling and simulation. Journal of computational chemistry 38, 1114–1124, doi:10.1002/jcc.24660 (2017).

33 Lee, J. et al. CHARMM-GUI supports the Amber force fields. The Journal of chemical physics 153, 35103, doi:10.1063/5.0012280 (2020).

34 Tian, C. et al. ff19SB: Amino-Acid-Specific Protein Backbone Parameters Trained against Quantum Mechanics Energy Surfaces in Solution. J Chem Theory Comput 16, 528–552, doi:10.1021/acs.jctc.9b00591 (2020).

35 Lee, J. et al. CHARMM-GUI supports the Amber force fields. J Chem Phys 153, 035103, doi:10.1063/5.0012280 (2020).

36 He, X., Man, V. H., Yang, W., Lee, T. S. & Wang, J. A fast and high-quality charge model for the next generation general AMBER force field. J Chem Phys 153, 114502, doi:10.1063/5.0019056 (2020).

37 Roe, D. R. & Cheatham III, T. E. PTRAJ and CPPTRAJ: software for processing and analysis of molecular dynamics trajectory data. Journal of chemical theory and computation 9, 3084–3095 (2013).

38 Harrigan, M. P. et al. MSMBuilder: Statistical Models for Biomolecular Dynamics. Biophysical journal 112, 10–15 (2017).

