## supplement data for "Molecular recognition of two endogenous hormones by the human parathyroid hormone receptor-1"

### Supplementary Materials

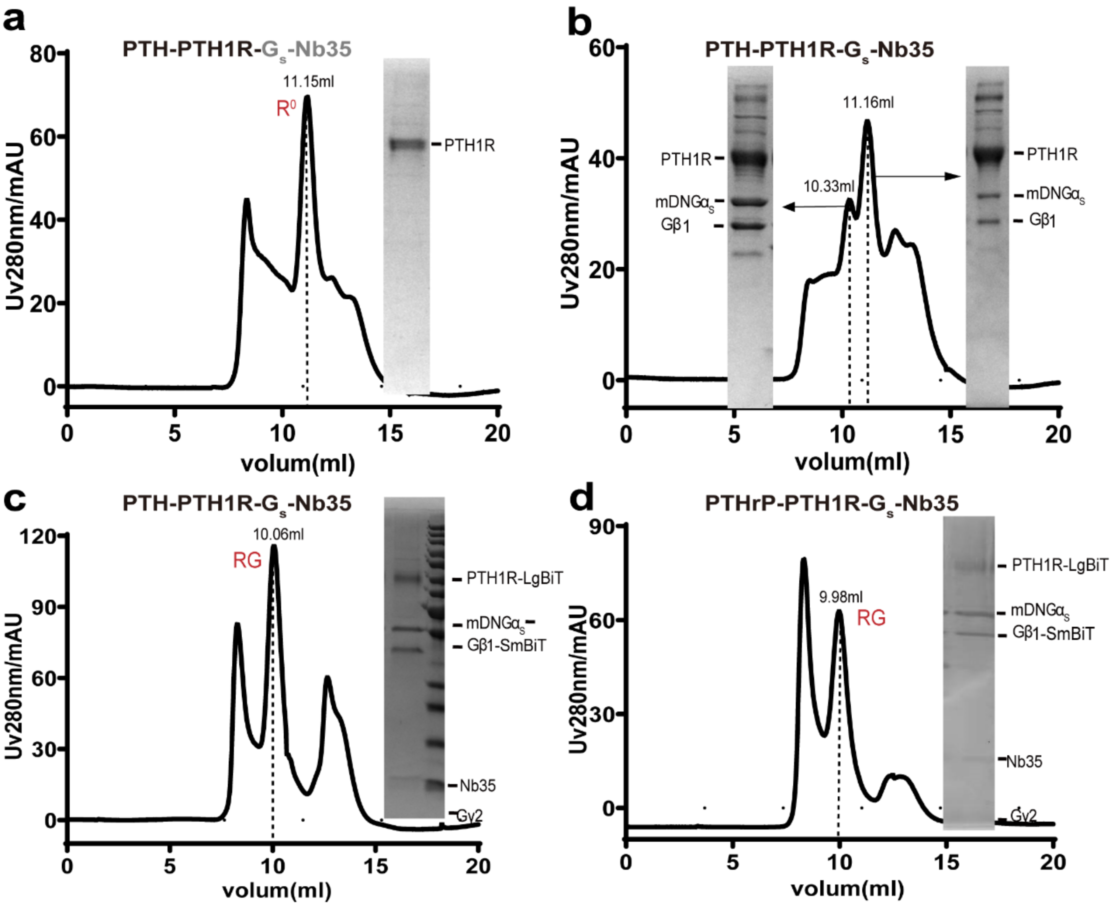


**Supplementary Fig. 1 Purification and characterization of the PTH-PTH1R-$G_s$ and PTHrP-PTH1R-$G_s$ complexes. (a)** The size-exclusion chromatography elution profile on Superdex200 Increase 10/300GL (left panel) and SDS-PAGE analysis (right panel) of the PTH-PTH1R ($R^0$) complex, PTH1R co-expression with DNG$\alpha_s$, Gβ1 and Gγ2, and purified with PTH, but no forming PTH-PTH1R-$G_s$ complex. **(b)** The size-exclusion chromatography elution profile on Superdex200 Increase 10/300GL (left panel) and SDS-PAGE analysis (right panel) of the PTH-PTH1R ($R^0$ and RG) complex, PTH1R co-expression with mDNG$\alpha_s$, Gβ1 and Gγ2, forming two states complexes peak. **(c-d)** The size-exclusion chromatography elution profile on Superdex200 Increase 10/300GL (left panel) and SDS-PAGE analysis (right panel) of the PTH-PTH1R-$G_s$ and PTHrP-PTH1R-$G_s$ complex (RG), which were adapted PTH1R-LgBiT to co-expression with mDNG$\alpha_s$, G$\beta$1-peptide 86 and G$\gamma$2.

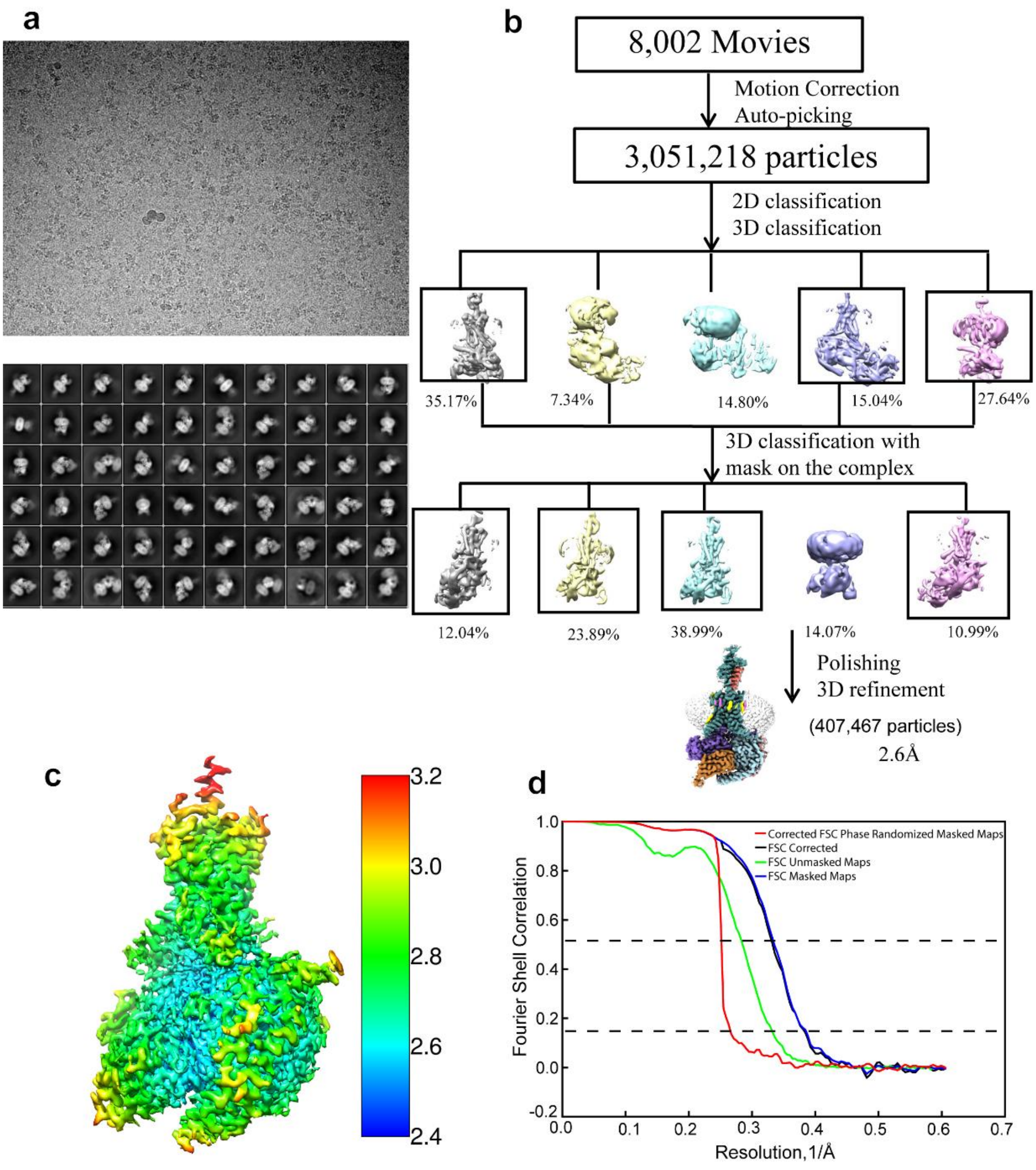


**Supplementary Fig. 2 Flowchart of cryo-EM data analysis of the PTH-PTH1R-G$_s$ complex. (a)** Representative cryo-EM micrographs of the PTH-PTH1R-G$_s$ complex and representative 2D class averages showing distinct secondary structure features from different views. **(b)** Flowchart of cryo-EM data analysis. **(c)** Cryo-EM map of the PTH-PTH1R-G$_s$ complex, colored by local resolution (Å) calculated using RELION. **(d)** "Gold-standard" FSC curve of the PTH-PTH1R-G$_s$ complex.

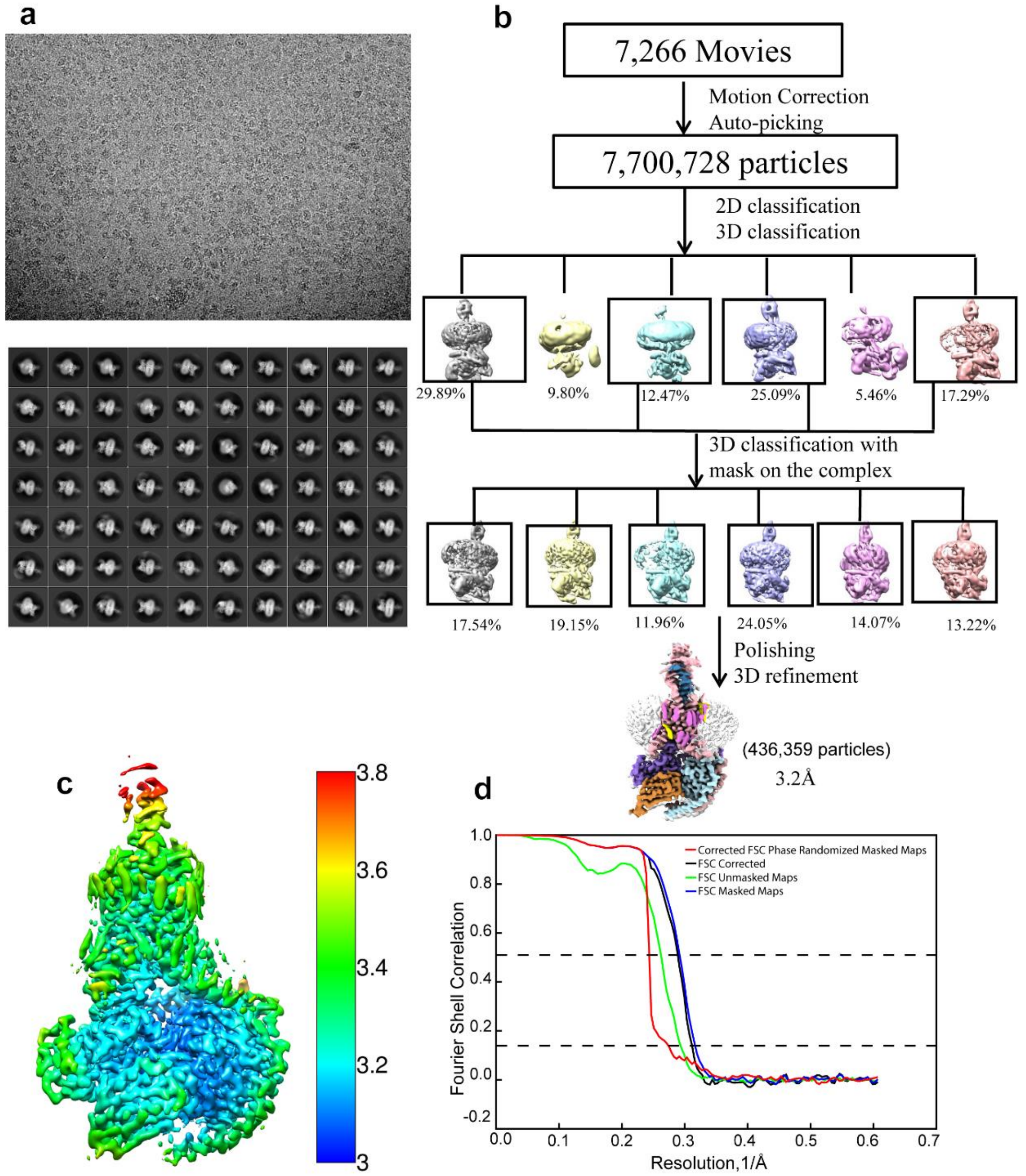


**Supplementary Fig. 3 Flowchart of cryo-EM data analysis of the PTHrP-PTH1R-$G_s$ complex.** (**a**) Representative cryo-EM micrographs of the PTHrP-PTH1R-$G_s$ complex and representative 2D class averages showing distinct secondary structure features from different views. **(b)** Flowchart of cryo-EM data analysis. **(c)** Cryo-EM map of the PTHrP-PTH1R-$G_s$ complex, colored by local resolution (Å) calculated using RELION. **(d)** “Gold-standard” FSC curve of the PTHrP-PTH1R-$G_s$ complex.

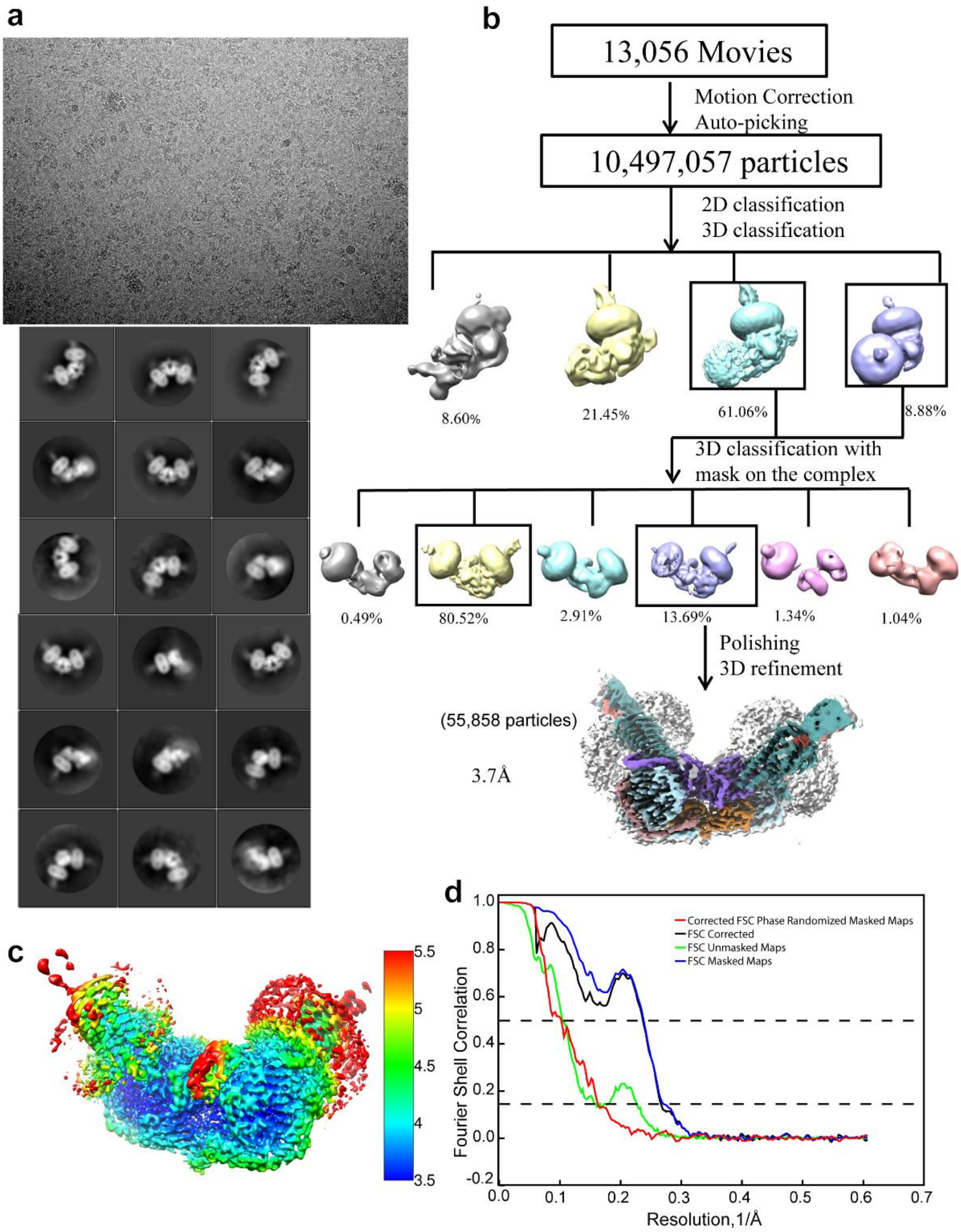


**Supplementary Fig. 4 Flowchart of cryo-EM data analysis of the PTHrP-PTH1R-$G_s$ complex dimer.** (**a**) Representative cryo-EM micrographs of the PTH-PTH1R-$G_s$ complex dimer and representative 2D class averages showing distinct secondary structure features from different views. **(b)** Flowchart of cryo-EM data analysis. **(c)** Cryo-EM map of the PTH-PTH1R-$G_s$ complex dimer, colored by local resolution (Å) calculated using RELION. **(d)** "Gold-standard" FSC curve of the PTH-PTH1R-$G_s$ complex dimer.

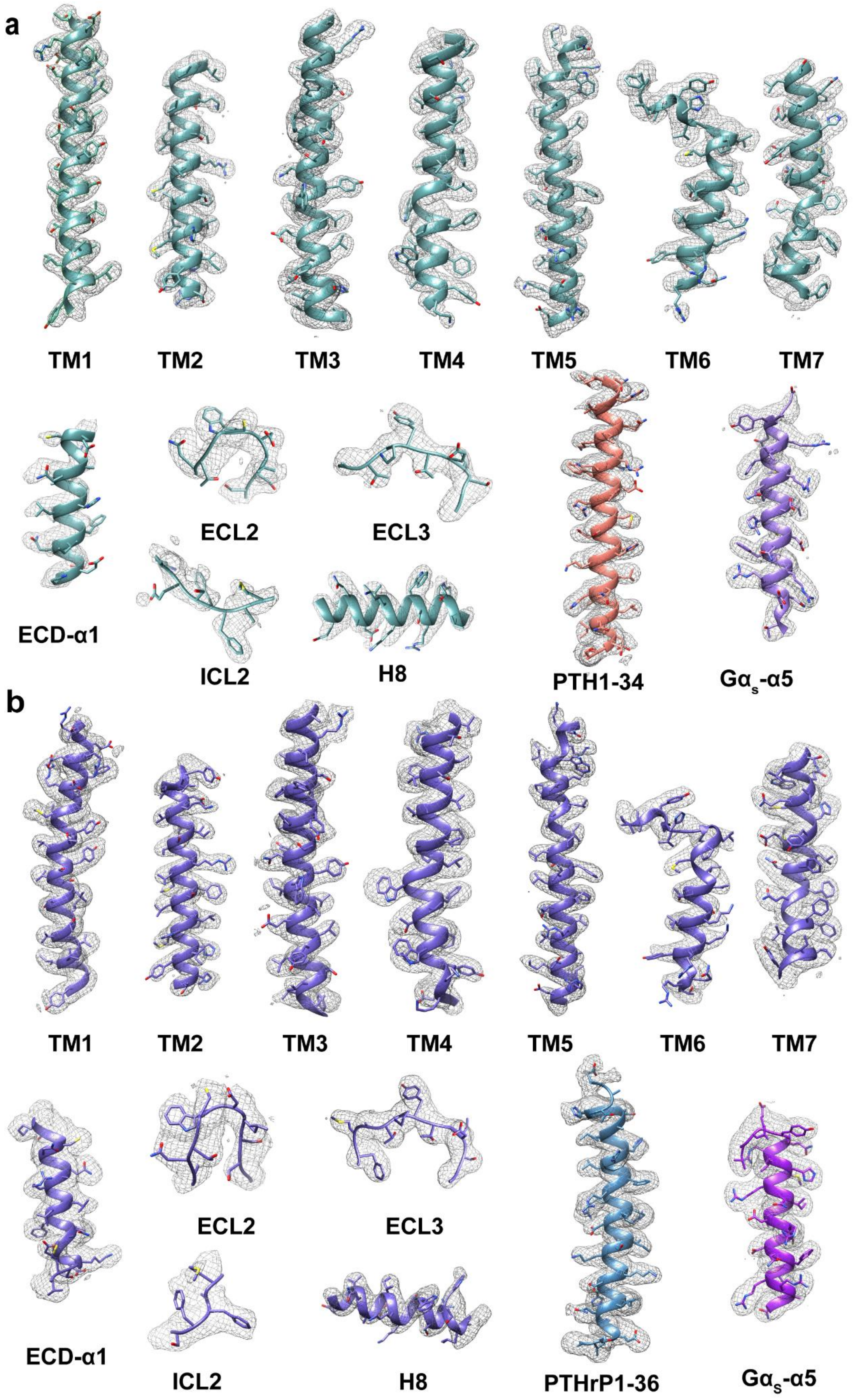
a
TM1
TM2
TM3
TM4
TM5
TM6
TM7
ECD-α1
ECL2
ECL3
ICL2
H8
PTH1-34
Gαs-α5
b
TM1
TM2
TM3
TM4
TM5
TM6
TM7
ECD-α1
ECL2
ECL3
ICL2
H8
PTHrP1-36
Gαs-α5

**Supplementary Fig. 5 Cryo-EM density maps of the ligand-PTH1R-$G_s$ protein structures. (a)** Cryo-EM density map and the model of the PTH-PTH1R-$G_s$ structure are shown for all transmembrane helices, ECD-α1 helix, ECL2, ECL3, ICL2 and helix 8 of PTH1R, PTH, and $G\alpha_s$-α5 helix. The model is shown in stick representation. **(b)** Cryo-EM density map and the model of the PTHrP-PTH1R-$G_s$ structure are shown for all transmembrane helices, ECD-α1 helix, ECL2, ECL3, ICL2 and helix 8 of PTH1R, PTHrP, and $G\alpha_s$-α5 helix. The model is shown in stick representation.

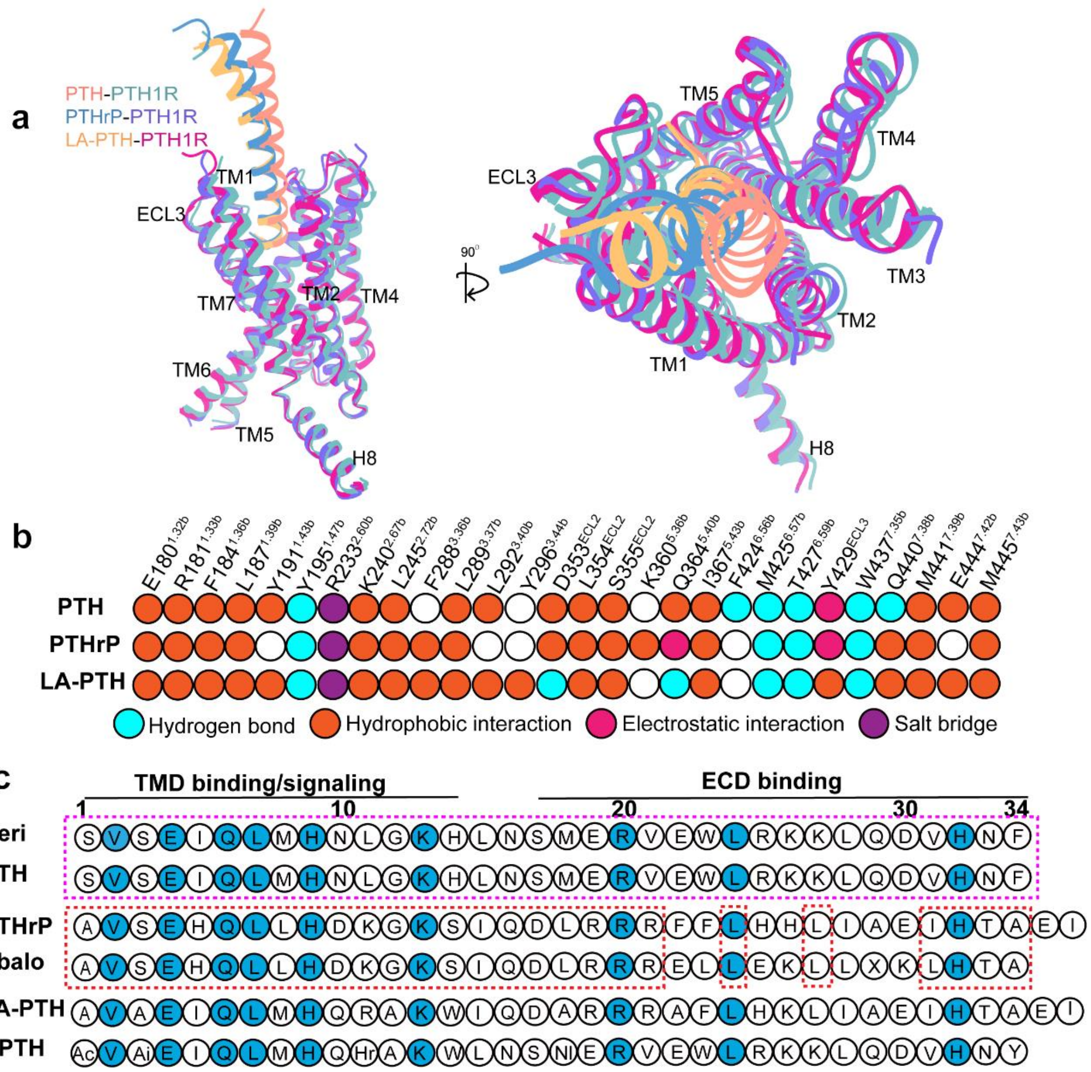


**Supplementary Fig. 6 Structural comparison of PTH1R-TMD bound by PTH, PTHrP and LA-PTH. (a)** Structural comparison of PTH1R-$G_s$ complexes bound by different ligands. Receptor ECD and G protein are omitted for clarity. **(b)** Comparison of the interactions of the PTH, PTHrP and LA-PTH N terminus with the PTH1R TMD. Receptor residues within 4.0 Å of PTH or PTHrP are listed. Color codes are listed on the bottom panel. Residues that show no interaction with ligands are displayed as white circles. **(c)** Sequence alignment of PTH1R agonists in single-letter code. Ac, aminocyclopentane-1- carboxylic acid; Ai, a-aminoisobutyric acid; Hr, homoarginine; Nl, norleucine; Teri: teriparatide; Abalo: abaloparatide.

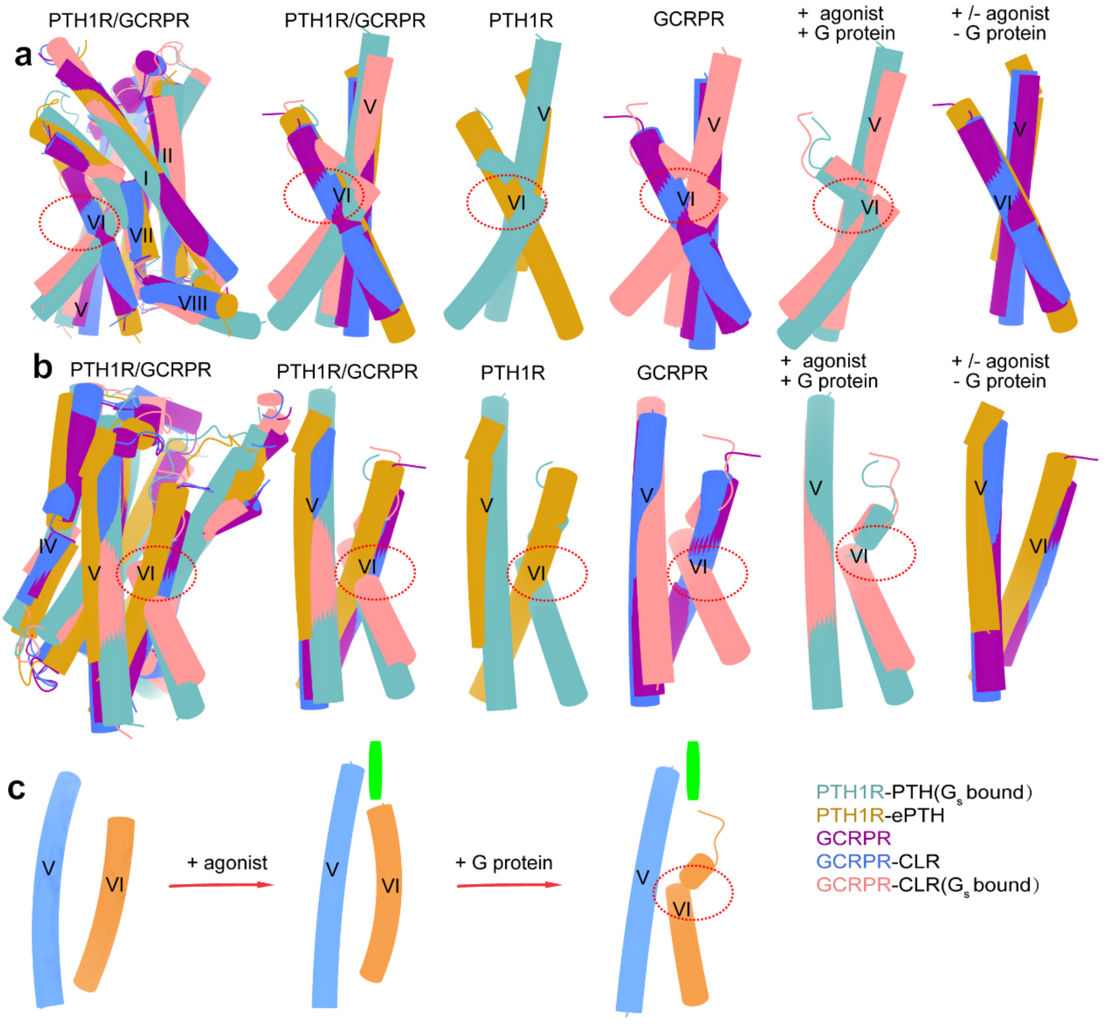


**Supplementary Fig. 7 Conformational changes during class B GPCRs activation.** **(a-b)** Different side views of superposition of PTH34-PTH1R-$G_s$, ePTH-PTH1R (PDB: 6FJ3), GCRPR-Ramp1 (PDB: 7KNT), CLR-GCRPR-Ramp1 (PDB: 7KNU) and CLR-GCRPR-Ramp1-$G_s$ (PDB: 6E3Y). Except for receptor TMD, others are omitted for clarity. **(c)** Schematic representation of TM6 conformational change during class B GPCRs activation. Left: inactive class B GPCRs (no agonist bound), middle: agonist bound class B GPCRs, but in the absence of G protein, and right: agonist bound class B GPCRs in G protein-bound state.

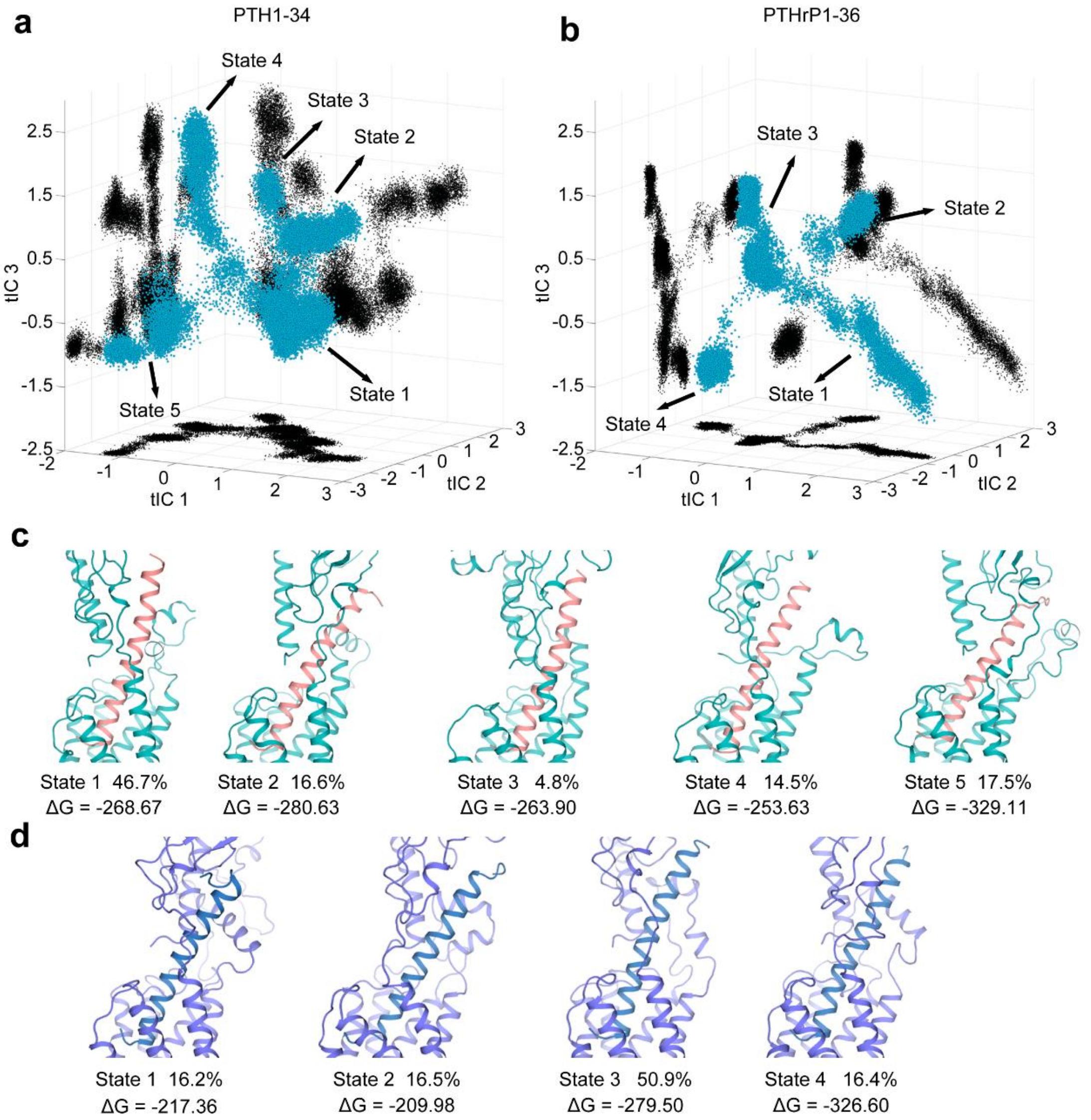


**Supplementary Fig. 8 Representative states analyzed at the interacting interface between the two endogenous hormones and PTH1R. (a-b)** The free energy landscape of PTH1-34 (a) and PTHrP (b) systems in 400 ns × 6 molecular dynamics (MD) simulations. The first three time-lagged independent component (tIC) 1, 2, and 3 were constructed according to the contacts between residue pairs of ligand and PTH1R to project the interactions onto the low-dimensional space. The corresponding macrostates are labeled by arrows. **(c-d)** The representative structure of each PTH1-34 **(c)** and PTHrP1-36 **(d)** macrostate with their proportion and binding free energy estimated by Molecular Mechanics Generalized Born Surface Area (MMGBSA). PTHrP1-36 has states 1 and 2 with high binding free energy values in states 1 and 2 (ΔG = -217.36 and -209.98 kcal/mol, respectively), which is worse than other states.

That phenomenon may explain why PTH1-34 has a greater capacity to bind to $R^0$ than PTHrP1-36.

**Supplementary Table 1. Cryo-EM data collection, refinement and validation statistics.**

| | PTH-PTH1R-$G_s$-complex | PTHrP-PTH1R-$G_s$-complex | Dimer |
|---|---|---|---|
| **Data collection and processing** | | | |
| Magnification | 105000 | 105000 | 105000 |
| Voltage (kV) | 300 | 300 | 300 |
| Electron exposure (e–/Å2) | 50 | 50 | 50 |
| Defocus range (μm) | -1.2 to -1.8 | -1.5 to -1.8 | -1.2 to -1.8 |
| Pixel size (Å) | 0.824 | 0.824 | 0.824 |
| Symmetry imposed | C1 | C1 | C1 |
| Initial particle images (no.) | 3,051,218 | 7,700,728 | 5,533,956 |
| Final particle images (no.) | 407,467 | 436,359 | 55,858 |
| Map resolution (Å) | | | |
| FSC threshold | 0.143 | 0.143 | 0.143 |
| Map resolution (Å) | 2.62 | 3.25 | 3.76 |
| Map sharpening B factor (Å2) | -69.26 | -85.21 | -75.59 |
| Refinement | | | |
| Initial model used (PDB code) | 6NBF/3C4M | 6NBF/3H3G | 6NBF/3C4M |
| Model resolution (Å) | 2.9 | 3.4 | 4.0 |
| FSC threshold | 0.5 | 0.5 | 0.5 |
| Model-Map CC (mask) | 0.79 | 0.80 | 0.66 |
| Model composition | | | |
| Non-hydrogen atoms | 9268 | 9312 | 36774 |
| Protein residues | 1160 | 1166 | 2320 |
| B factors (Å2) | | | |
| Protein | 60.66 | 91.30 | 59.4 |
| R.m.s. deviations | | | |
| Bond lengths (Å) | 0.002 | 0.003 | 0.002 |
| Bond angles (Å) | 0.431 | 0.542 | 0.499(2) |
| Validation | | | |
| MolProbity score | 1.38 | 1.61 | 1.51 |
| Clash score | 6.48 | 8.12 | 5.72 |
| Rotamer outliers (%) | 0.1 | 0.2 | 0.25 |
| Ramachandran plot | | | |
| Favored (%) | 97.89 | 97.03 | 96.38 |
| Allowed (%) | 2.02 | 2.79 | 3.03 |
| Disallowed (%) | 0.09 | 0.17 | 0.18 |

**Supplementary Table 2. Effects of PTH and PTHrP bind to PTH1R WT and mutants.**

| Mutant | PTH | | PTHrP | |
|---|---|---|---|---|
| | $pEC_{50}$ ± S.E.M. | $E_{max}$ ± S.E.M. (% WT) | $pEC_{50}$ ± S.E.M. | $E_{max}$ ± S.E.M. (% WT) |
| WT | 11.28 ± 0.03 | 99.97 ± 073 | 10.94 ± 0.08 | 100.25 ± 1.84 |
| M32A | 10.93 ± 0.09 | 99.69 ± 1.89 | 9.79 ± 0.05* | 99.63 ± 1.78 |
| K34A | 11.10 ± 0.07 | 99.36 ± 1.73 | 10.84 ± 0.05 | 99.27 ± 1.43 |
| E35A | 11.00 ± 0.10 | 99.21 ± 2.35 | 9.8 ± 0.05* | 98.94 ± 1.53 |
| Y136A | 10.84 ± 0.09 | 98.87 ± 1.81 | 10.29 ± 0.06 | 98.02 ± 1.92 |
| D137A | 9.70± 0.13* | 99.67 ± 3.36 | 7.40 ± 0.14*** | 105.77 ± 8.85 |
| Y167A | 9.38 ± 0.10** | 99.11 ± 2.72 | 7.99 ± 0.09*** | 100.17 ± 3.77 |
| E180A | 11.22 ± 0.11 | 97.37 ± 2.24 | 10.46 ± 0.06 | 97.78 ± 1.49 |
| R181A | 11.58 ± 0.09 | 97.93 ± 1.58 | 10.90± 0.07 | 97.52 ±1.56 |
| Y195F | 9.19 ± 0.08*** | 98.60 ± 2.51 | 7.74 ± 0.10*** | 100.84 ± 4.65 |
| R233A | 8.46 ± 0.17*** | 97.83 ± 5.20 | N. A. | N. A. |
| L292A | 8.23 ± 1.47*** | 18.29 ± 5.34 | 7.04 ± 1.02*** | 7.08 ± 7.35 |
| D353A | 11.28 ± 0.06 | 97.94 ± 1.30 | 9.80 ± 0.06* | 96.17 ± 1.76 |
| Q364A | 10.95 ± 0.07 | 98.69 ± 1.44 | 9.82 ± 0.07* | 96.90 ± 1.88 |
| M425A | 10.03 ± 0.10 | 97.45 ± 2.24 | 9.49 ± 0.07** | 94.31 ± 1.82 |
| Y429A | 10.50 ± 0.10 | 99.08 ± 2.01 | 10.18 ± 0.08 | 97.78 ± 2.01 |
| T430A | 11.07 ± 0.07 | 98.90 ± 1.68 | 9.95 ± 0.05 | 96.42 ± 1.46 |
| W437A | 9.77 ± 0.13* | 98.93 ± 3.42 | 6.95 ± 0.10*** | 109.68 ± 7.42 |
| Q440A | 10.41 ± 0.08 | 99.80 ± 2.15 | 9.78 ± 0.06* | 98.15 ± 1.73 |
| M441A | 10.51 ± 0.09 | 100.25 ± 2.31 | 7.86 ± 0.09*** | 101.48 ± 4.34 |
| E444A | 10.90 ± 0.09 | 101.22 ± 1.98 | 10.16 ± 0.09 | 98.63± 2.68 |
| M445A | 11.35 ± 0.08 | 100.43 ± 1.61 | 9.85 ± 0.06* | 100.02 ± 1.89 |

cAMP accumulation assay was performed in wild-type (WT) and single-point mutated PTH1R expressing in HEK293T cells. cAMP accumulation data were analyzed using a three-parameter logistic equation to determine $pEC_{50}$ and Emax values. $E_{max}$ values for mutants are showed as a percentage of the WT. Data were generated and graphed as means ± S.E.M. of three independent experiments (n = 3) performed in quadruplicate. One-way ANOVA were used to determine statistical difference (*P< 0.05, **P< 0.01, ***P< 0.001).
